# Diversity and pathobiology of an ilarvirus unexpectedly detected in diverse host plants and in global sequencing data

**DOI:** 10.1101/2022.12.15.520526

**Authors:** Mark Paul Selda Rivarez, Chantal Faure, Laurence Svanella-Dumas, Anja Pecman, Magda Tušek-Žnidaric, Deborah Schönegger, Kris De Jonghe, Arnaud Blouin, Sebastien Massart, Maja Ravnikar, Denis Kutnjak, Armelle Marais, Thierry Candresse

**Affiliations:** Department of Biotechnology and Systems Biology, National Institute of Biology, 1000 Ljubljana, Slovenia; Université de Bordeaux, INRAE, UMR 1332 Biologie du Fruit et Pathologie, 33882 Villenave d’Ornon, France; Plant Sciences Unit, Flanders Research Institute for Agriculture, Fisheries and Food, 9820 Merelbeke, Belgium; Plant Pathology Laboratory, TERRA-Gembloux Agro-Bio Tech, University of Liège, 5030 Gembloux, Belgium

**Author notes:** corresponding authors: M.P.S. Rivarez and T. Candresse. College of Agriculture and Agri-industries, Caraga State University, Ampayon, Butuan City, 8600 Agusan del Norte, Philippines (Mark Paul Selda Rivarez, Arnaud Blouin). Plant Protection Department, Agroscope, 1260 Nyon, Switzerland.

**Keywords:** *Ilarvirus*, *Solanaceae*, Serratus, virus diversity, phylogenetics, histopathology, virion morphology, symptomatology, virus transmission, pollen

## Abstract

High-throughput sequencing (HTS) and sequence mining tools revolutionized virus detection and discovery in recent years and implementing them with classical plant virology techniques results to a powerful approach to characterize viruses. An example of a virus discovered through HTS is Solanum nigrum ilarvirus 1 (SnIV1) (family *Bromoviridae*), which was recently reported in various solanaceous plants from France, Slovenia, Greece, and South Africa. It was likewise detected in grapevines (*Vitaceae*) and several *Fabaceae* and *Rosaceae* plant species. Such a very diverse host association is atypical for ilarviruses, thus warranted further investigation. In this study, modern and classical virological tools were combined to accelerate the characterization of SnIV1. Through HTS-based virome surveys, mining of sequence read archive datasets, and literature search, SnIV1 was further identified from diverse plant and non-plant sources globally. SnIV1 isolates showed relatively low variability compared to other phylogenetically related ilarviruses. Phylogenetic analyses showed a distinct basal clade of isolates from Europe, while the rest formed clades of mixed geographic origin. Furthermore, systemic infection of SnIV1 in *Solanum villosum* and its mechanical and graft transmissibility to solanaceous species were demonstrated. Near identical SnIV1 genomes from the inoculum (*S. villosum*) and inoculated *Nicotiana benthamiana* were sequenced, thus partially fulfilling Koch’s postulates. SnIV1 was shown to be seed-transmitted and potentially pollen-borne, has spherical virions, and possibly induces histopathological changes in infected *N. benthamiana* leaf tissues. Overall, this study provided information to better understand the diversity, distribution, and pathobiology of SnIV1, but whether it could emerge as a destructive pathogen remains uncertain.

**Funding:**

1. EU Horizon 2020 Marie Skłodowska-Curie Actions Innovative Training Network (H2020 MSCA-ITN) project no. GA 813542
2. Administration of the Republic of Slovenia for Food Safety, Veterinary Sector and Plant Protection and Slovenian Research Agency (ARRS) funding no. P4-0165, P4-0407, J4-4553
3. Balik Scientist Program (Republic Act 11035) of the Department of Science and Technology– Philippine Council for Agriculture, Aquatic, and Natural Resources Research and Development (DOST–PCAARRD), Republic of the Philippines
4. The Belgian FPS Health Food Chain Safety and Environment under Project RT18/3 SEVIPLANT

## INTRODUCTION

The combination of classical virology techniques and modern high-throughput sequencing (HTS) and bioinformatics tools provides a powerful approach to detect, identify and characterize viruses and monitor changes in their populations even before they emerge and cause problems (Maclot et al. 2020; McLeish et al. 2021; Kumar et al. 2022). The COVID-19 pandemic and the persistent risks posed by plant and animal virus diseases to our food supply (Morens et al. 2020; Ristaino et al. 2021; Meurens et al. 2021) have increased interest in viromic surveys of ecosystems and data-driven virus discovery (Carroll et al. 2018; Lauber and Seitz 2022). This led to a recent surge in the discovery of viruses and other virus- or viroid-like agents from various studies (Gregory et al. 2019; Edgar et al. 2022; Lee et al. 2022; Mifsud et al. 2022; Neri et al. 2022; Rivarez et al. 2022; Zayed et al. 2022). As a result, hundreds of thousands of putative novel viruses remain uncharacterized due to the astounding amount of experimental work their characterization would require. For plant virologists, it is an immense task to uncover the biological properties of newly identified plant viruses and to systematically assess their possible economic and biosecurity risks (Massart et al. 2017; Hou et al. 2020; Rivarez et al. 2021; Lebas et al. 2022).

Some recent studies combined classical and modern tools and techniques to characterize recently discovered plant viruses. For instance, the biological characterization of an emerging pathogen of tomato, Physostegia chlorotic mottle alphanucleorhabdovirus (family *Rhabdoviridae*), was significantly accelerated through an international collaboration driven by HTS data (Temple et al. 2022). Recently, mining of vast amounts of *Arabidopsis thaliana* publicly available sequence read archive (SRA) datasets uncovered a novel comovirus (family *Secoviridae*), Arabidopsis latent virus 1, which was demonstrated to be mechanically and seed transmitted, but causes no symptoms in *A. thaliana*, (Verhoeven et al. 2022). A recent study on *Prunus*-associated luteoviruses (family *Luteoviridae*) also uncovered a new luteovirus through a search of SRA datasets (Khalili et al. 2022). Many HTS-based discoveries of crop and non-crop viruses have also been reported (Gaafar et al. 2020; Rivarez et al. 2022; Ma et al. 2019; Xu et al. 2017), but only a small subset of these studies biologically characterized the identified viruses (Hou et al. 2020; Rivarez et al. 2021). Thus, there is a very wide knowledge gap concerning the biological properties of newly discovered plant viruses, which limits the evaluation of their economic and phytosanitary risks.

This study focused on Solanum nigrum ilarvirus 1 (SnIV1), a virus that was recently associated with wild, weedy or cultivated species in the *Solanaceae* family and for which only marginal biological information is available. SnIV1 was detected in *Solanaceae* crop and non-crop plants from France (*Solanum nigrum* and *S. lycopersicum*), Slovenia (*Physalis* sp.), South Africa (*S. chenopodiodes*), and Greece (*Capsicum annuum*) (Ma et al. 2020; Rivarez et al. 2022; Mahlanza et al. 2022; Orfanidou et al. 2022). The recent report from Greece also demonstrated the infectivity of SnIV1 in *Nicotiana benthamiana* and peppers (Orfanidou et al. 2022). Interestingly, SnIV1 was concurrently reported under different names [*i.e.*, grapevine-associated ilarvirus (GaIV), surrounding legume-associated ilarvirus (sLaIV), and Erysiphe necator-associated ilar-like virus 1 (EnaIV1)] in viromic studies involving plants from different botanical families. These studies detected SnIV1 sequences in legume (*Fabaceae*) plants from Germany (Gaafar et al. 2020) and grapevines (*Vitaceae*) from Italy and Spain (Chiapello et al. 2019, 2020). Recently, SnIV1 has also been reported in *Rosaceae* fruit trees such as peaches (*Prunus persica*) from the USA (Dias et al. 2022) and apricots (*Prunus armeniaca*) from South Africa (Bester and Maree 2022). Such a diverse list of host plants is unusual for a member of the *Ilarvirus* genus, which usually have host ranges limited to species of the same family, and prompted the current study.

SnIV1 belongs to the genus *Ilarvirus*, which is the largest genus in the family *Bromoviridae* with 22 recognized species, most of which are known plant viruses, with some causing significant economic losses (Simkovich et al. 2021). Ilarviruses are transmitted by vegetative propagation (Uyemoto 1992) and several ilarviruses are known to be pollen and/or seed transmitted (Mink 1993; Card et al. 2007). Their transmission could likewise be facilitated by thrips or by pollinators such as bees (*Apis* spp.), usually through infected pollen (Bristow and Martin 1999). Ilarviruses pose persistent threats to fruit production such as for *Prunus* species (Pallas et al. 2012; Khalili et al. 2022) or blackberry (*Rubus fruticosus*) (Poudel et al. 2014). In recent years, several ilarviruses were reported to cause problems in tomato production in the USA, such as tomato necrotic streak virus (TomNSV) (Badillo-Vargas et al. 2016; Adkins et al. 2015) and tomato necrotic spot virus (ToNSV) (Bratsch et al. 2019, 2018). Other ilarviruses also known to be emerging or endemic pathogens of tomatoes include Parietaria mottle virus (PMoV) that is mostly found in Europe (Aparicio et al. 2018), tobacco streak virus (TSV) that is endemic in tomato and several wild *Solanaceae* species (Sharman et al. 2015; Hanč spinach latent virus (SpLV) that was associated with symptomatic tomatoes (Vargas-Asencio et al. 2013). In the *Ilarvirus* genus subgroup 1, ToNSV, PMoV, and TSV are the most closely related viruses to SnIV1 (Ma et al. 2020; Bratsch et al. 2019).

In this study, aside from modern HTS and sequence mining approaches, classical techniques such as experimental host range tests, extensive geographical surveys, histopathological observations, diversity and phylogenetic analyses, and subsequent virus detection using RT-PCR tests were implemented. The general aim was to assess the global diversity and distribution and to characterize some biological properties of SnIV1 and, specifically, to answer the following questions: (1) Can the geographic distribution and associated hosts of SnIV1 be expanded using an HTS-based viromic survey of various plant species and by searching relevant SRA datasets? (2) What is the level of genetic diversity of SnIV1 relative to other related species? (3) Do isolates of SnIV1 show some form of phylogenetic structuring? (4) Can SnIV1 infect a range of experimental host plants? (5) What are the potential routes of transmission of SnIV1? (6) What is the virion morphology of SnIV1? and (7) Can some histopathological changes be associated with SnIV1 infection?

## MATERIALS AND METHODS

### Plant samples

The following plant species were collected in the INRAE Bordeaux research Center (Villenave d’Ornon, France), tested for SnIV1 infection, and/or sequenced in the surveys of this study: (1) *Solanum villosum* that underwent Nanopore sequencing and RT-PCR testing, (2) *S. nigrum* that underwent RT-PCR testing, (3) *Vitis vinifera* cultivar (cv.) Sauvignon, (4) *V. vinifera* cv. Ugni Blanc, and (5) *Daucus carota* subspecies (subsp.) *carota* that underwent Illumina sequencing and RT-PCR testing. Samples of *S. melongena* and *S. tuberosum* were similarly obtained from selected farms in Belgium and were submitted for Illumina sequencing and underwent further RT-PCR testing. Details on how RNA was extracted and sequenced are presented below.

Following positive RT-PCR tests for SnIV1, two *S. villosum* plants were uprooted from the field, cleaned, pruned, and introduced in an insect-proof greenhouse. Two SnIV1-positive grapevine samples (*V. vinifera* cv. Sauvignon) were introduced in the same greenhouse by preparing and transplanting cleaned stem cuttings from each plant. Both samples were maintained for three months in the greenhouse before retesting for SnIV1 and utilizing them in subsequent experiments as described below.

**Nucleic acid extraction methods for RT-PCR assays and HTS.** Different methods were used for RNA extraction from field samples, greenhouse-introduced plants, or inoculated test plants prior to RT-PCR testing and/or HTS on Illumina platforms.

For total RNA extractions performed in France, a previously described method (Foissac et al. 2005) was used for leaves, stems, seeds, fruits, roots, pollen, and floral parts of *S. villosum* and for the pollen of *S. nigrum* prior to RT-PCR testing. Likewise, this protocol was used to extract total RNAs individually from inoculated test plants prior to RT-PCR testing. The Spectrum^TM^ Total Plant RNA kit (Sigma-Aldrich, Saint-Quentin-Fallavier, France) was used, following the kit instructions, to extract total RNA from grapevine leaves, petioles, bark, and phloem scrapings for RT-PCR testing. A previously described total RNA extraction protocol (Svanella-Dumas et al. 2022) was used for individual grapevine leaf tissues (for cv. Ugni Blanc) or phloem scrapings (cv. Sauvignon) prior to HTS. Double-stranded RNA (dsRNA) from 45 pools of carrot plants (50 plants each) were purified as previously described protocol (Ma et al. 2020) prior to HTS (Schönegger et al., in preparation).

Virion-associated nucleic acid (VANA) were purified from pools of 50 individual samples each for *S. melongena* and *S. tuberosum* from Belgium prior to HTS as previously described (Palanga et al. 2016; Hammond et al. 2020).

Prior to RT-PCR assays of inoculated test plants from Slovenia, RNeasy^TM^ Plant Mini Kit (Qiagen, USA) was used to extract total RNAs from all samples following the kit instructions.

### Nanopore sequencing

A CTAB-based protocol (Chang et al. 1993) was used to extract total RNAs from SnIV1-infected *S. villosum* that served as inoculum and from inoculated *N. benthamiana* prior to Nanopore sequencing. Total RNAs extracted were then treated with DNase using the TURBO DNA-free™ Kit (Thermo Fisher Scientific, USA). RNA quality and quantity were checked prior to sequencing using an Epoch^TM^ microplate spectrophotometer (BioTek, Agilent, USA) and a QuBit fluorometer (Thermo Fisher Scientific, USA). Nanopore sequencing was performed as previously described (Rivarez et al. 2022; Pecman et al. 2022). Briefly, total RNAs were depleted of ribosomal RNA using the RiboMinus^TM^ Plant Kit (Thermo Fisher Scientific, USA) and polyadenylated using *E. coli* Poly(A) polymerase (New England Biolabs, UK). Sequencing libraries were prepared using the PCR-cDNA barcoding kit (catalog no. SQK-PCB109, version 10Oct2019, Oxford Nanopore Technologies (ONT), UK), prior to sequencing using the ONT MinION platform (ONT, UK) with base-calling following a previously described workflow (Pecman et al. 2022).

### RT-PCR assays

Oligonucleotide primers specific for SnIV1 RNA 3 segment or tomato betanucleorhabdovirus 2 (TBRV2, a virus detected in mixed infection with SnIV1 in *S. villosum*) (Table 1) were used in RT-PCR tests conducted in Slovenia using the OneStep^TM^ RT-PCR kit (Qiagen, USA) as previously described (Rivarez et al. 2022). Additional SnIV1 primers targeting RNA 1 and RNA 3 segments were designed using OligoCalc (Kibbe 2007) and used in two-step RT-PCR reactions as previously described (Marais et al. 2014), to test for SnIV1 in different plant samples from France. In each RT-PCR assay, RNA extracts from SnlV1-positive samples were used as positive control, RNA extraction control and/or healthy plants as negative controls, and no template (water only) as blank control.

**TABLE 1.**
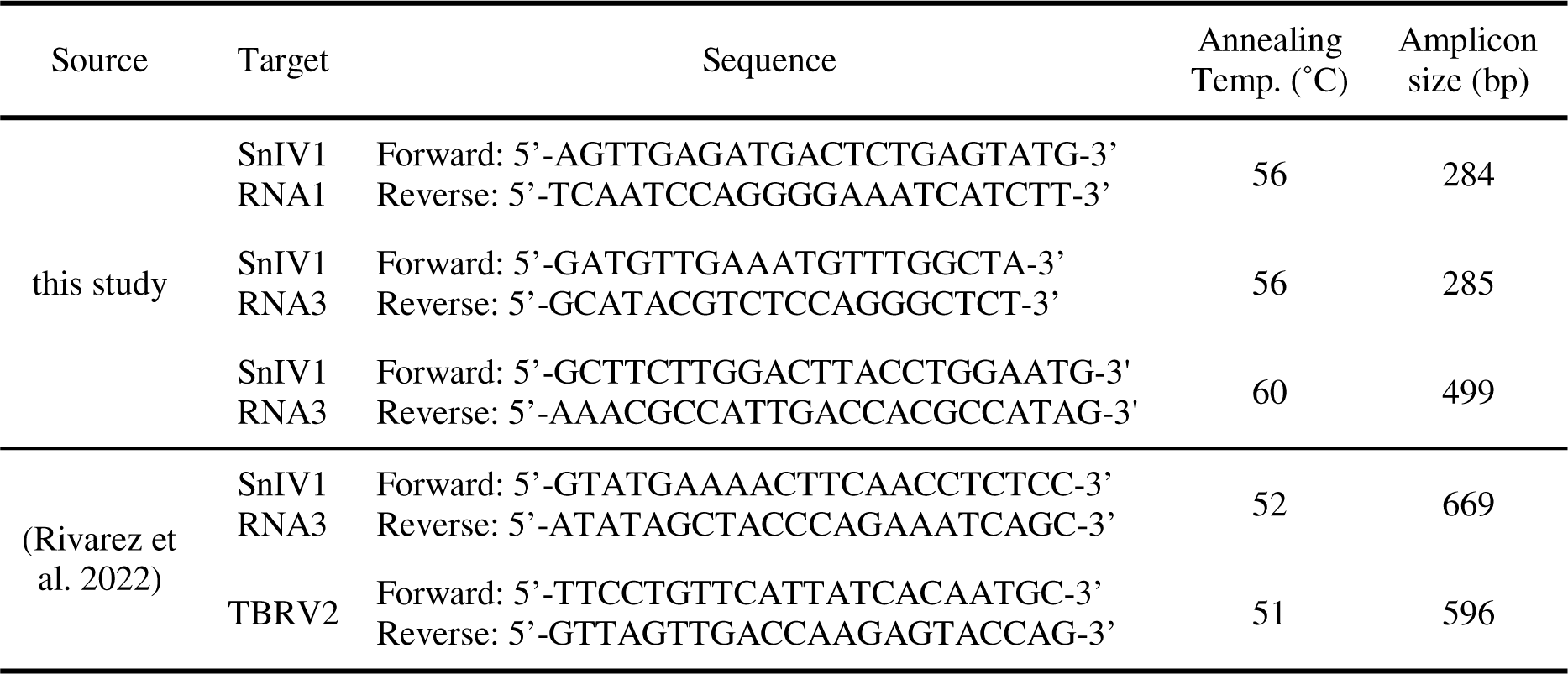
RT-PCR primer pairs used for the detection of SnIV1 and TBRV2.

### Search for SnIV1 sequences in databases and the literature

Publicly available databases and the literature were searched for SnIV1 sequences to obtain additional SnIV1 genomes. Firstly, NCBI GenBank database release version (r.v.) 250 that includes nucleotide, whole genome shotgun (WGS), transcriptome shotgun assembly (TSA) sequences, *etc.* (Sayers et al. 2022) was searched using the BLASTn algorithm (Altschul et al. 1997), and with SnIV1 genomic segments (GenBank accession number (a.n.) OL472060-OL472062) as queries. The search was performed on July 22, 2022. Percent identity and percent query coverage values were both set at greater than 90% as threshold to select significant BLASTn hits.

Secondly, a search in global metagenomes, metatranscriptomes, or metaviromes was performed through a palmID search in Serratus (https://serratus.io/palmid) (Edgar et al. 2022), which analyzed and annotated in a database (palmDB) around 5.7 million global SRA datasets deposited as of January 2021. The SnIV1 RNA 2 sequence (a.n. OL472061) which contains the RNA-dependent RNA polymerase (RdRp) domain needed for the palmID search was used as query. A 100% RdRp ‘palmprint’ (Babaian and Edgar 2021) amino acid identity and an E-value less than 8.4x10^-74^ were used as thresholds to identify significant palmID hits.

Third, published literature was searched using Google Scholar (https://scholar.google.com/) with the queries ‘Solanum nigrum ilarvirus 1’, ‘grapevine-associated ilarvirus’, or ‘surrounding legume-associated ilarvirus’, which are the first three virus names proposed for SnIV1. The search was performed on August 3, 2022. SnIV1 sequences that were not yet available in GenBank r.v. 250 were requested from the authors of the studies that identified the virus prior to their release in NCBI or as official publication.

### Assembly of SnIV1 genomes

A previously described bioinformatic pipeline was used for the reference-guided assembly of SnIV1 genomes from HTS data in this study (Pecman et al. 2017; Rivarez et al. 2022). Using the pipeline that was run in CLC Genomics Workbench (GWB) version (v.) 20 (Qiagen, USA), barcodes were removed from raw reads that were previously screened based on phred quality scores. Virus and virus-like reads and contigs were initially identified by mapping reads and contigs to the virus RefSeq database r.v. 212 (Sayers et al. 2022) and by viral domain searches in contigs against the pFam v. 33 (Mistry et al. 2021). SnIV1 consensus genomes were reconstructed by mapping reads to the genome of a Slovenian isolate (a.n. OL472060-OL472062), with percent identity and genome coverage threshold set at ≥ Mapping profiles were visually inspected and read mapping depth (r.m.d.) or on average, the number of times each locus in a reference genome is covered by mapped reads was noted. The number of mapped reads, percent genome covered, and the presence of a complete set of open reading frames (ORFs) were noted, in reference to some recommendations for the detection of virus genomes in metagenomic data (Roux et al. 2019; Simmonds et al. 2017). Genomes assembled from SRA datasets were deposited in GenBank as third party annotations.

In parallel efforts, a previously described method was used to *de novo* assemble and annotate contigs from HTS data obtained from Belgian samples (Buzkan et al. 2019). When needed, contigs representing SnIV1 genome segments were extended by iterative mapping of reads as implemented in Geneious Prime v. 2022.1.1 (Dotmatics, USA).

A customized workflow (Pecman et al. 2022) was used to assemble SnIV1 genomes from ONT MinION sequencing data of *S. villosum* (inoculum) and inoculated *N. benthamiana*. The workflow was used for quality screening, barcode trimming, demultiplexing, visualization, and *de novo* assembly. Minimap2 v. 2.24 (Li 2018) as implemented in Geneious Prime v. 2022.1.1 (Dotmatics, USA) was used for mapping reads or contigs to viral RefSeq r.v. 212 (Sayers et al. 2022) and SnIV1 genome (a.n. OL472060-OL472062).

### Multiple sequence alignments and recombination detection analyses.

To examine the molecular diversity of SnIV1, nucleotide (nt) sequences were aligned using MUSCLE (Edgar 2004) as implemented in MEGA X (Kumar et al. 2018). To compare SnIV1 diversity with that of other ilarviruses, sequences of isolates of tobacco streak virus (TSV) and Parietaria mottle virus (PMoV) were retrieved from GenBank r.v. 250 and aligned similarly for further analyses.

Prior to diversity and phylogenetic analyses, possible recombination events among the SnIV1, TSV, or PMoV sequences were checked using RDP v. 4 (Martin et al. 2015). A recombination event was considered significant if it has a *p*-val<10^-4^ in at least four of the methods used (RDP, GENECONV, Bootscan, Maxchi, Chimaera, SiSscan, PhylPro, LARD, 3Seq) (de Klerk et al. 2022; Stewart et al. 2014). Recombinant sequences were removed and unaligned ends for each genome segment of the remaining isolates manually trimmed.

### Nucleotide diversity and genetic variation analyses

The movement protein (MP) and coat protein (CP) ORFs from the RNA 3 segment of each viral isolate were concatenated into a contiguous sequence. Since most isolates have full RNA 3 sequence, their MP and CP were concatenated to achieve uniformity and retain coding information for all isolates in the subsequent analyses. The RNA 1, RNA 2, and concatenated RNA 3 alignments were then used separately to perform pairwise identity and nucleotide diversity analyses. For each genome segment, pairwise identities were calculated using SDT v. 1.2 (Muhire et al. 2014).

Genome-wide polymorphisms were detected and nucleotide diversities [pi (π)] and molecular genetic variation [theta (θ), based on and finite sites model] (Subramanian 2016) were calculated with DnaSP v. 6 (Rozas et al. 2017) using a sliding windows of 30 bases and a step size of 3. Overall genetic distances were calculated in MEGA X (Kumar et al. 2018) using the same set of alignments.

### Phylogenetic analyses

Maximum likelihood phylogenetic analyses were used to examine the clustering of SnIV1 isolates from diverse sources. Multiple sequence alignments described above were used as input for the analyses performed in MEGA X (Kumar et al. 2018). The most suitable substitution model was selected based on the Bayesian information criteria, and the analyses were performed with 1,000 bootstrap replicates. iToL v. 6.4 (Letunic and Bork 2021) was used to visualize and annotate the resulting phylogenetic trees.

### Disinfection of plant tissues and seeds

To remove possible surface contaminants, including pollen grains possibly carrying SnIV1, plant tissues (including seeds) were surface disinfected prior to RNA extraction. This was done on plant samples from France including the greenhouse-introduced *S. villosum* and grapevine (cv. Sauvignon) tissues, as well as leaves of inoculated test plants. Disinfection was done by soaking the tissues in a 5% sodium hypochlorite solution for 10 min with intermittent agitation, followed by six washes in sterile water with blot-drying in between. The sodium hypochlorite solution and water were replaced for every new tissue fragment being disinfected and washed. After the washes, disinfected plant tissues were air-dried for at least 15 min before proceeding with RNA extraction.

### Preparation of floral parts and pollen for RNA extraction

Individual floral parts and pollen were tested for the presence of SnIV1. Ten flowers from the greenhouse-introduced, SnIV1-infected *S. villosum* plant (described above) were collected. Pedicels, sepals, pistils, stamens, and petals were dissected and separately pooled prior to RNA extraction.

Pollen grains were collected from the same *S. villosum* plant described above and from *S. nigrum* inoculated by approach grafting (see details below). Briefly, ripe stamens were separated from flowers and vortexed in sterile water to liberate and suspend pollen grains. Stamens were then removed, and purity and integrity of pollen grains verified under the Eclipse Ni-U (Nikon, Japan) microscope with a dark field condenser in reflection mode. Pollen grains suspended in sterile water were briefly ground using a sterile plastic pestle suitable for 1.5 ml Eppendorf tubes before proceeding with RNA extraction.

### Mechanical transmission tests.

The greenhouse-introduced SnIV1-infected *S. villosum* was used as the inoculum source for the mechanical inoculations. Twenty plants per host plant species with three plants for each species kept as mock-inoculated control were used as test plants. The inoculum was prepared by homogenizing 1.0 g infected tissue with 10 ml phosphate buffer (0.02 M, pH 7.8, supplemented with 0.112 g sodium diethyldithiocarbamatetrihydrate β-mercaptoethanol per 100 ml total volume). Activated charcoal powder (0.1 g per 10 ml inoculum) was added to the ice-cold inoculum which was then used to rub-inoculate plants using approximately 0.1 ml inoculum on the second and third youngest leaves of plants that was dusted with carborundum. Plants were maintained in an insect-proof greenhouse with temperature set at 20-24°C, with 16/8 h day/night cycle. Samples from individual plants or pooled equal amounts of uninoculated newly formed leaves from inoculated plants were tested for SnIV1 presence, up until 35 days post inoculation (dpi).

### Graft transmission tests.

The greenhouse-introduced, SnIV1-infected *S. villosum* was used in approach-grafting transmission experiments by using the healthy, greenhouse-grown *S. nigrum* plants as recipients (n=3). Briefly, about 2-3 cm vertical length of the epidermal-parenchymal layer was removed on one side of a young stem in both source and recipient plants. The exposed tissues were joined together and secured with a perforated adhesive tape.

For chip bud grafting, the same infected *S. villosum* plant was used to obtain 2-3 cm superficial tissue strip pieces from of a young stem, which were joined with the exposed internal tissues of a young stem of recipient *S. nigrum* plants (n=2). The joined tissues were again secured with a perforated adhesive tape. Both approach- and chip bud-grafted plants were maintained in greenhouse maintained at 20-24°C, with 16/8 h day/night cycle, before RT-PCR testing for SnIV1 at five weeks after grafting.

### Seed transmission tests.

Seeds were collected from a greenhouse-introduced SnIV1-infected *S. villosum* (n=73) and from two randomly selected *S. nigrum* and two *S. villosum* plants from the field (INRAE, Bordeaux, France), and surface disinfected as described above. A subsample of these seeds was also tested by RT-PCR to confirm SnIV1 infection prior to sowing. Seeds were sown in a soil tray and maintained in the greenhouse with conditions set at 20-24°C, with 16/8 h day/night cycle. RT-PCR testing of germinated seedlings for SnIV1 infection was performed at three and five weeks after sowing.

### Microscopic examination of SnIV1 virions and infected leaf tissues.

Leaves of the same age and size from mock-inoculated and SnIV1-infected *N. benthamiana* plants were sampled at 49 dpi. For negative staining, leaf tissue homogenates were prepared by macerating them in 1.5 ml Eppendorf tubes containing phosphate buffer (0.1 M, pH 7.0). The homogenates were applied to Formvar-coated, carbon-stabilized copper grids and negatively stained with 1% uranyl acetate (SPI Supplies, USA) in phosphate buffer (0.1 M, pH 7.0) before inspection using a Talos^TM^ transmission electron microscope (TEM) (ThermoFisher, USA).

For preparation of thin tissue sections for light microscopy, small pieces of the same leaves used for TEM observations were fixated in 3% glutaraldehyde in phosphate buffer (0.1 M, pH 7.0) for 16 h at 4 °C, which was followed by post-fixation in 1% osmium tetroxide in phosphate buffer (0.1 M, pH 7.0) and embedding in Agar 100 resin (Agar Scientific, UK). Semi-thin sections (0.6 µm) were cut with a Reichert Ultracut S ultramicrotome (Leica, Germany), stained with Azure II/methylene blue, and observed with an Axioskop 2 Plus microscope (Carl Zeiss, Germany).

## RESULTS

### SnlV1 genome sequences obtained from HTS of different plant species

SnIV1 genomes were sequenced and assembled in four different HTS experiments (Table 2). Nanopore long read sequencing of rRNA-depleted total RNA from *S. villosum* and from *N. benthamiana* yielded high coverage SnIV1 genomes. In this experiment, r.m.d. ranged from 29x to 15,841x with 100% genome coverage in all three genome segments. Full genome sequences of TBRV2 were also recovered from both plants and are 90-95% identical to TBRV2 genomes deposited in GenBank from a previous study (Rivarez et al. 2022) (Supplementary Fig. 1). This is the first detection of TBRV2 in a wild plant (*S. villosum*) and in France. However, TBRV2 was not identified from the HTS datasets of the other sequenced samples discussed below.

**TABLE 2.**
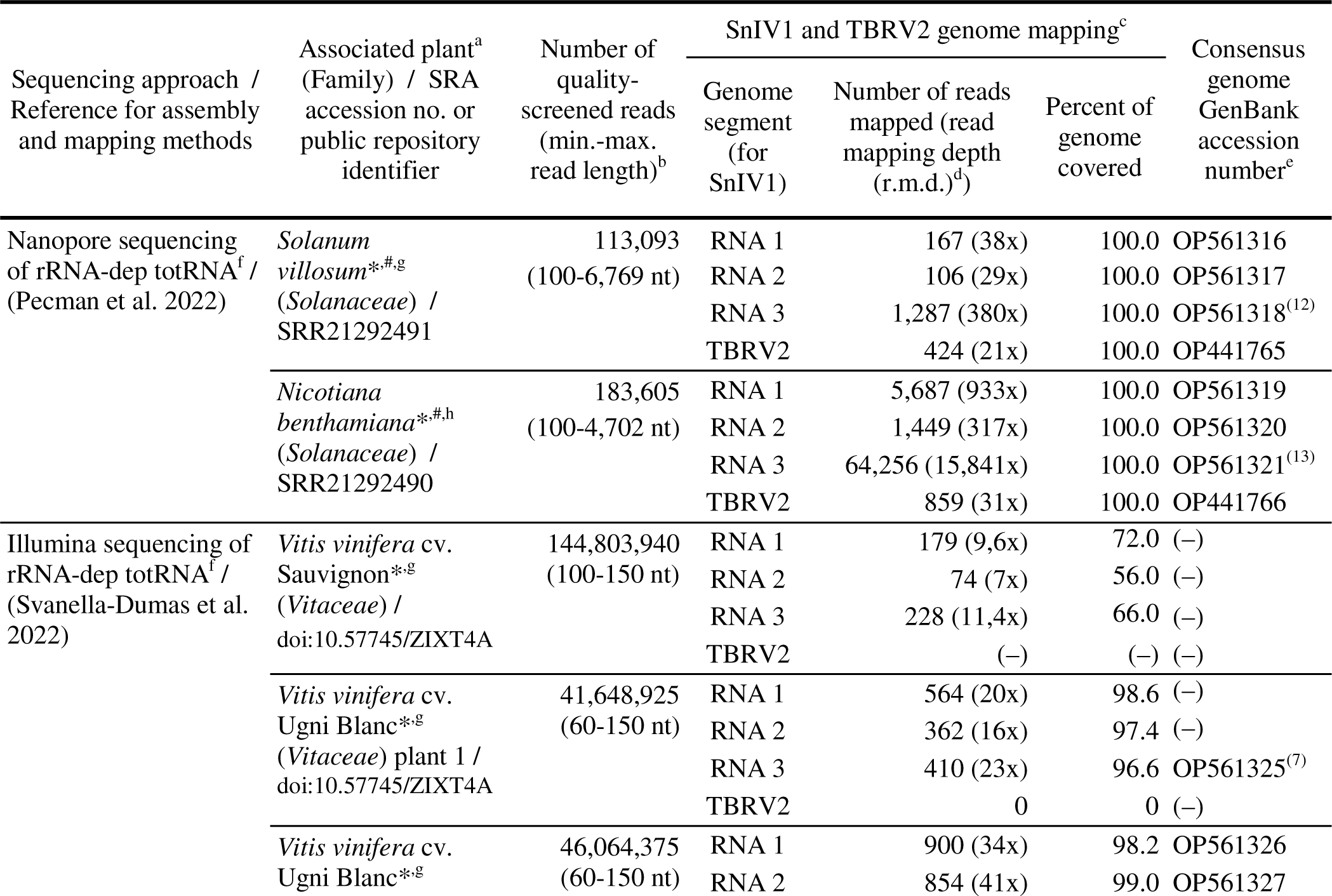
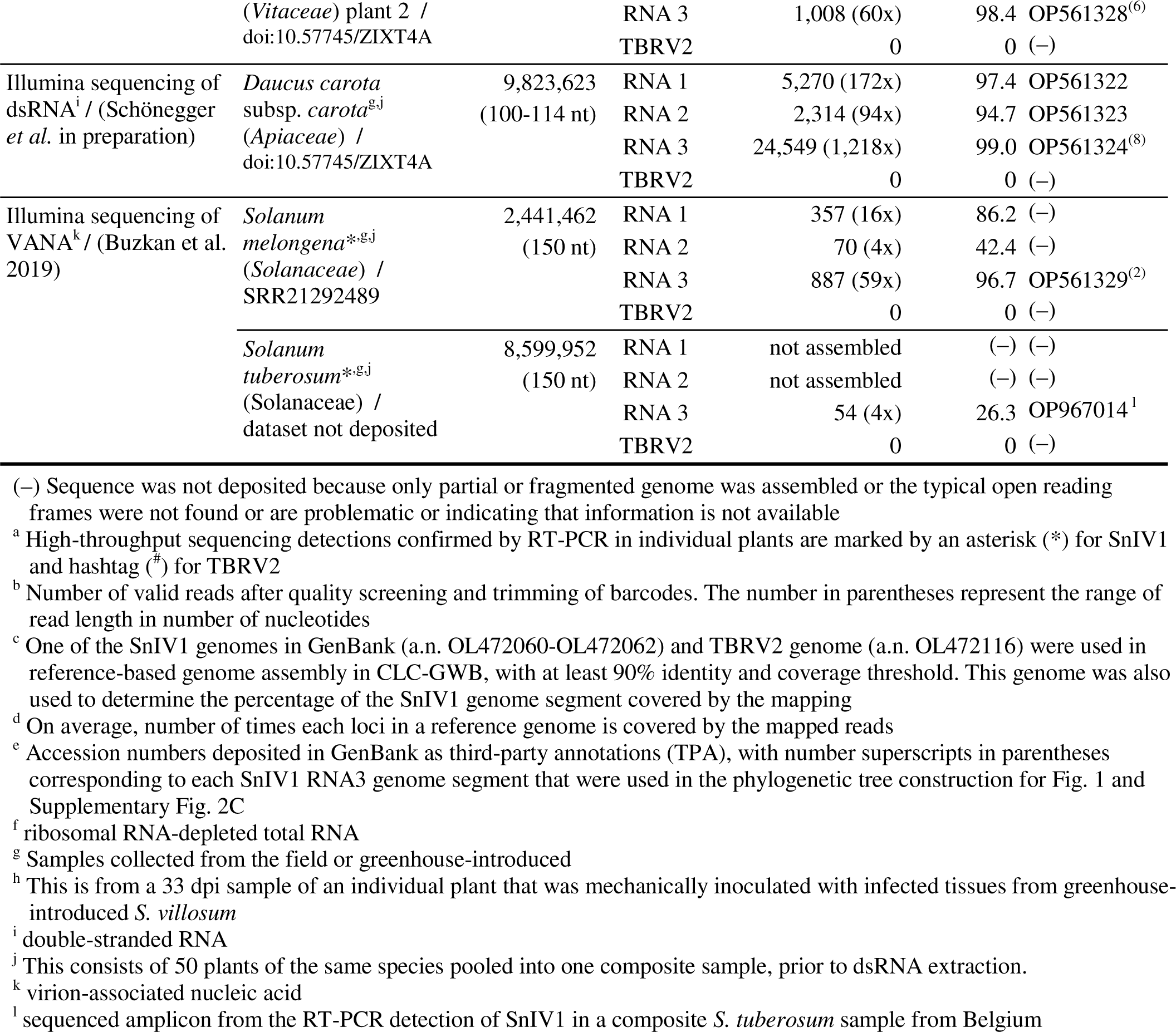
High-throughput sequencing of plant samples collected from field surveys and those collected from transmission experiments emphasizing the detection of SnIV1 sequences and its genome assembly.

Viromic studies using plants collected during the field surveys in France and Belgium yielded partial, near complete or complete genomes of SnIV1. Illumina sequencing of dsRNA from wild carrots yielded near complete SnIV1 genome segments with an average r.m.d. ranging between 94x and 1,218x for the three segments. Positive RT-PCR tests confirmed the presence of SnIV1 in the dsRNA extract of the pooled wild carrot samples.

HTS of a pool of five grapevine (cv. Sauvignon) phloem scrapings samples yielded 481 reads that mapped on SnIV1 genome segments. RT-PCR tests confirmed the presence of SnIV1 in two of the five grapevines. Stems of the positive grapevines were later cleaned and introduced in the greenhouse as cuttings and, after growing for several months, were used in the subsequent RT- PCR tests on different grapevine tissues (see below). Illumina short read sequencing of rRNA-depleted total RNA from two grapevine (cv. Ugni Blanc) plants yielded near complete SnIV1 genomes with average r.m.d. of 16x-60x. This detection was later confirmed by RT-PCR tests for both individual samples.

Illumina sequencing of VANAs from a pool of *S. melongena* and *S. tuberosum* samples collected in Belgium yielded partial genome of SnIV1, with only a near complete RNA 3 segment assembled from the *S. melongena* dataset. The detection of SnIV1 in pooled samples of both species was later confirmed with a positive RT-PCR test. The amplicon from *S. tuberosum* was Sanger sequenced and confirmed to be SnIV1.

RT-PCR detection of SnIV1 in different plant tissues.

The presence of SnIV1 was further investigated in different tissues of SnIV1-positive *S. villosum* and grapevines (cv. Sauvignon) that were introduced and grown for several months in the greenhouse. Three out of eight grown cuttings each from the two Sauvignon grapevines were sampled for bark, phloem, petiole, and newly grown leaf tissues. All tissues tested negative for SnIV1 at four and eight months post-introduction in the greenhouse (Table 3). However, leaves from the same grapevines collected in the field, before the plants were introduced in the greenhouse, had previously tested positive using RT-PCR.

**TABLE 3.**
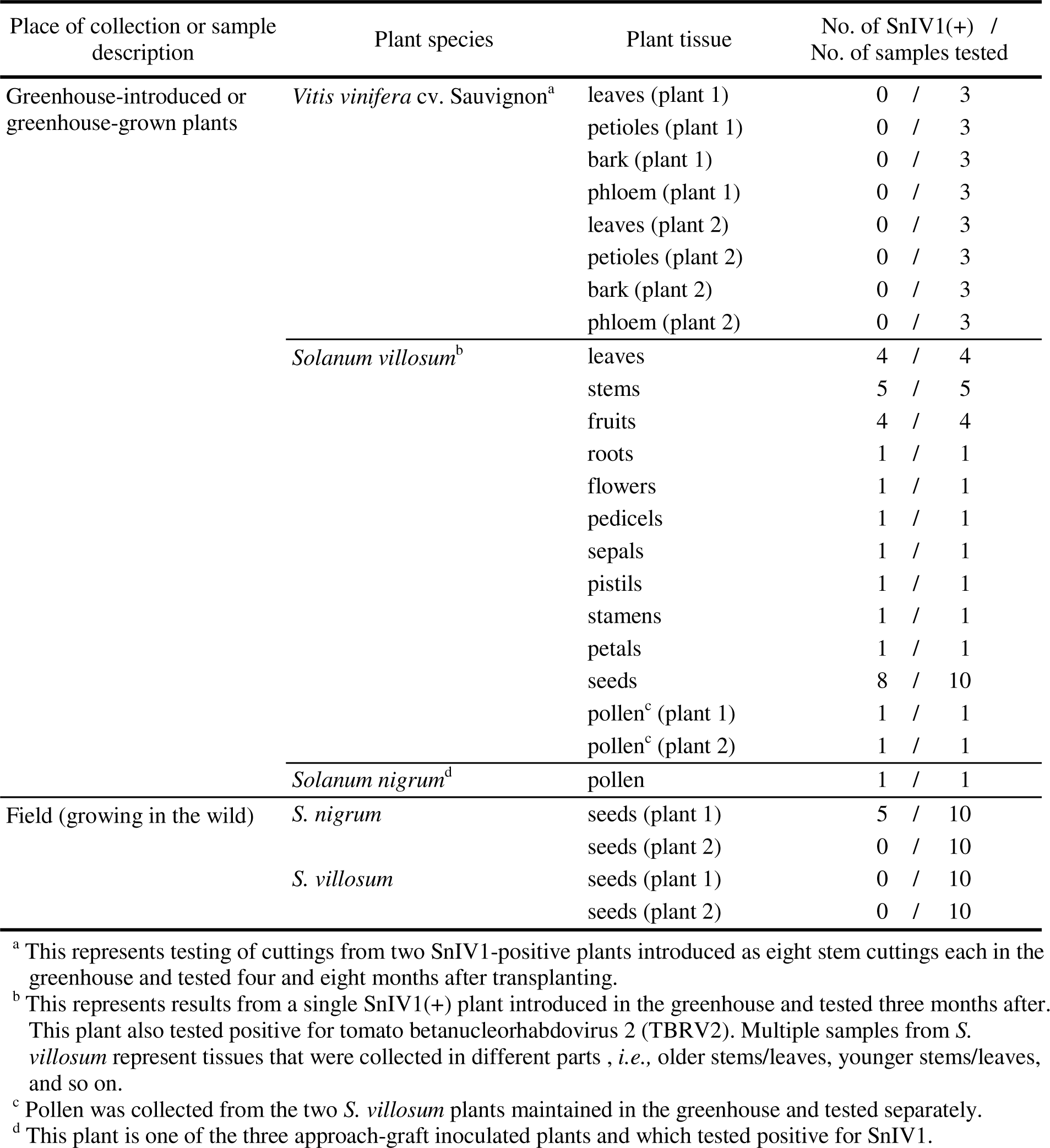
RT-PCR detection of SnIV1 in different tissues from greenhouse-introduced plants or from plants growing in the field. Place of collection or sample description

Disinfected tissues of one of the two *S. villosum* plants replanted in the greenhouse (without showing virus disease symptom) were also tested three months after introduction of the plant in the greenhouse. All *S. villosum* tissues that were surface disinfected and tested in pools (*i.e.*, leaf, stem, fruits, root, flowers, and seeds pools), as well as non-disinfected individual floral parts and pollen, tested positive for SnIV1.

The presence of SnIV1 was also evaluated in pollen from a graft-inoculated *S. nigrum* plant (see details below) maintained in the greenhouse and in seeds from randomly sampled wild *S. nigrum* and *S. villosum* collected from on the INRAE Bordeaux research center. Specifically, pollen collected from the graft-inoculated *S. nigrum* plant tested positive for SnIV1. RT-PCR tests of surface-disinfected individual seeds revealed the presence of SnIV1 in seeds from only one of the two randomly sampled *S. nigrum* plants from the field, while seeds from two *S. villosum* plants that were similarly processed tested negative for SnIV1.

### *In silico* detection of SnIV1 in databases and in the literature

Information on sequences and existing records of SnIV1 were collected to gain a comprehensive picture of its global diversity and geographic distribution (Table 4). Aside from SnIV1 genomes deposited in GenBank database r.v. 250, SnlV1 was also detected through BLASTn homology searches in a publicly available TSA of hop (*Humulus lupulus* var. *lupulus*, family *Cannabaceae*) from Japan (Natsume et al. 2015). So far, this is the only detection of SnIV1 sequences that are linked to samples from Asia and deposited in the GenBank database. Furthermore, search of SRA datasets through palmID in Serratus (Edgar et al. 2022) returned 80 datasets with RdRp ‘palmprint’ sequences that are 100% identical to that of SnIV1 (103 amino acid residues, E-value < 10^-74^).

**TABLE 4.**
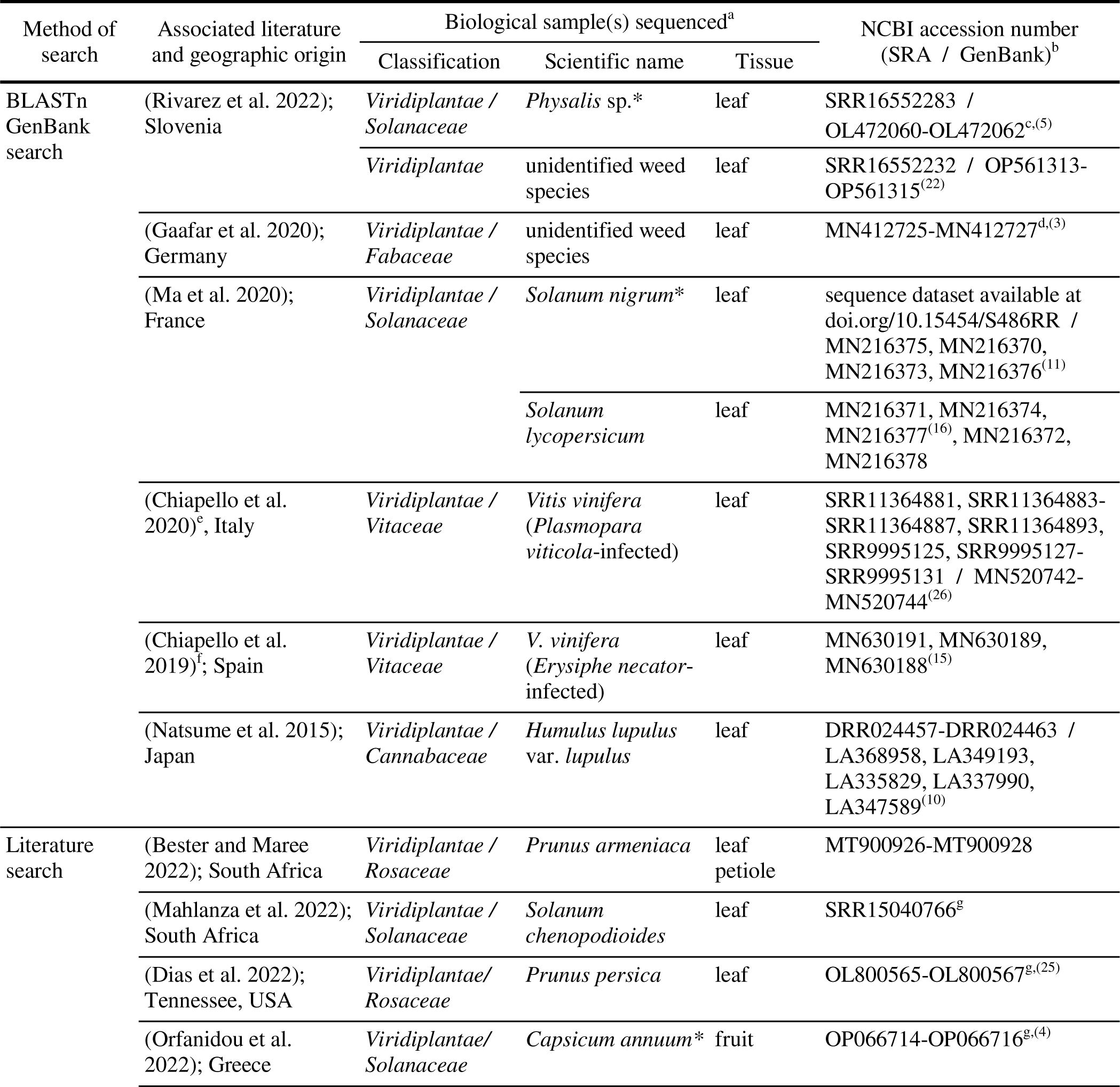
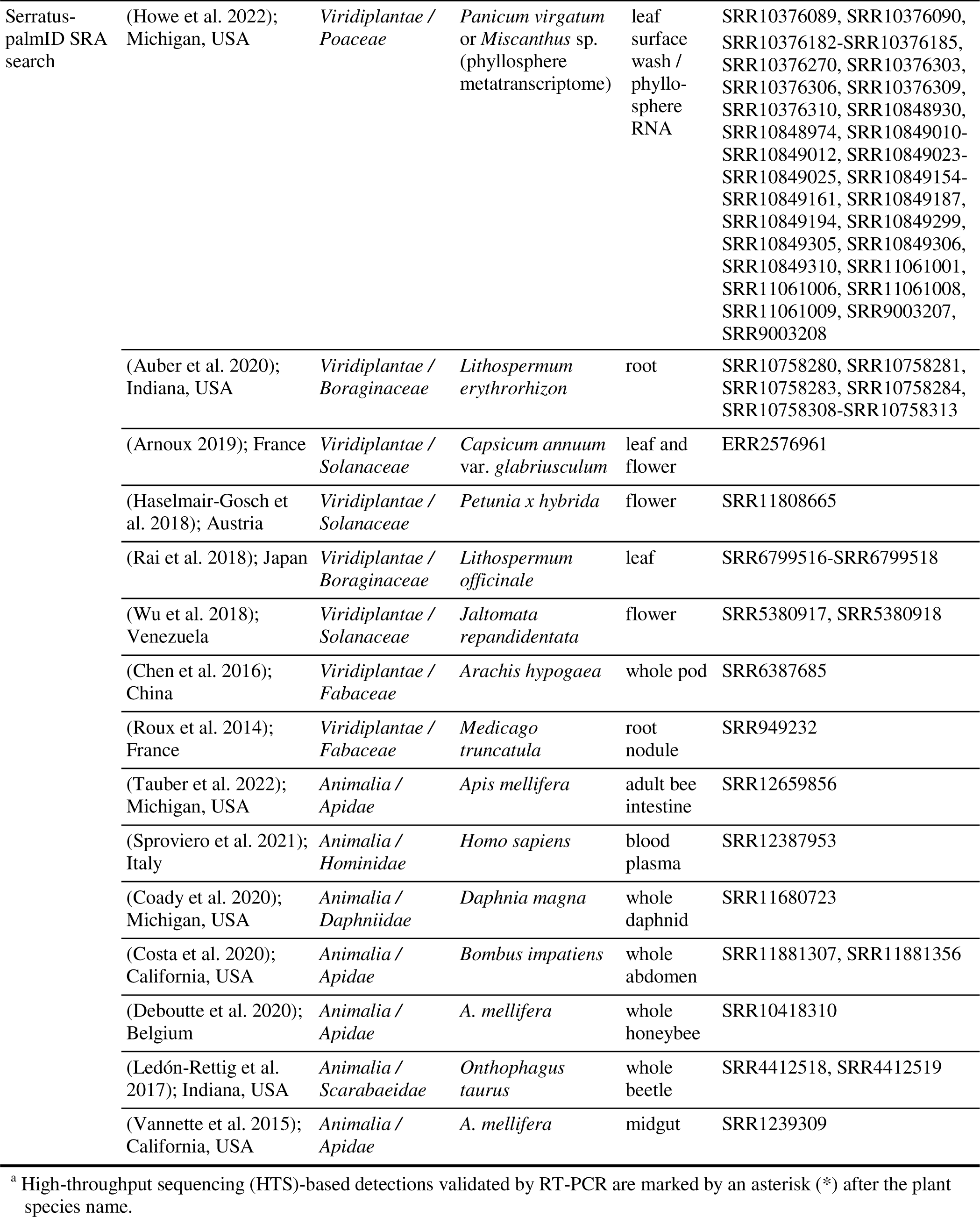
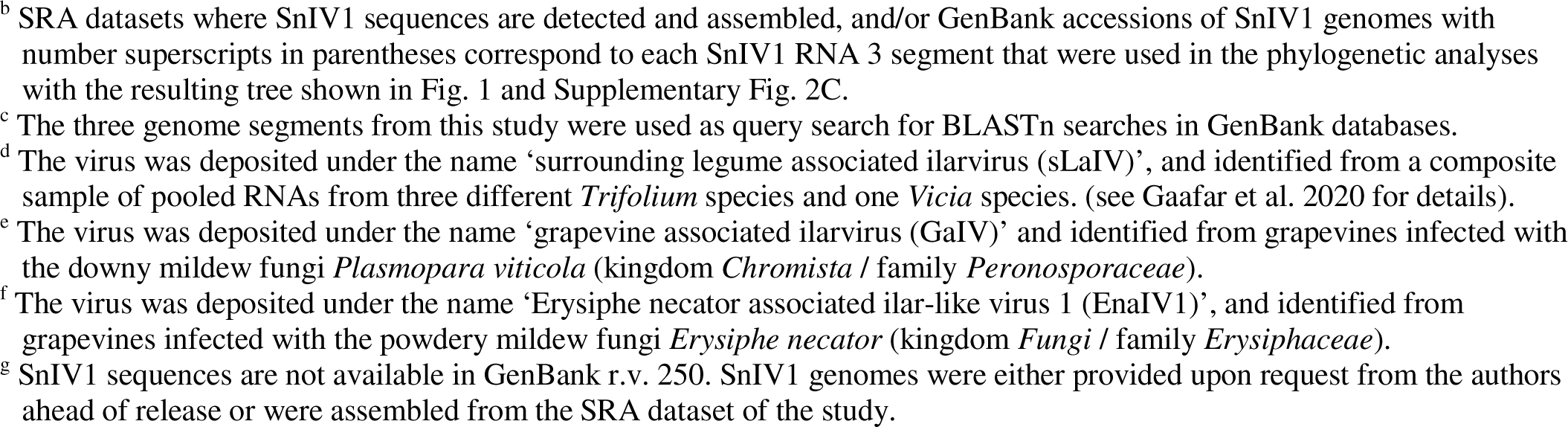
Detection of SnIV1 from *in silico* searches of public sequence databases, sequence read archive datasets, and reports from literature. All BLASTn hits are >90% identical to SnIV1 sequences (a.n. OL472060-OL472062) (E-value < 10^-4^), while Serratus-palmID hits are 100% identical to SnIV1 RdRp palmprint (E-value < 10^-74^).

These results included detections of SnIV1 in sequence datasets from China and the USA and several European countries.

A literature search identified recent studies that detected SnIV1 sequences that were not yet available in GenBank r.v. 250. This search identified four new plant viromic or disease etiology studies, including two that detected the virus in South Africa, which represent the first reports of SnIV1 in the African continent (Mahlanza et al. 2022; Bester and Maree 2022). The other two studies are viromic studies of peach in the USA (Dias et al. 2022), and an HTS study of symptomatic peppers (hybrid Arlequin F1) from Greece (Orfanidou et al. 2022). The study from Greece demonstrated the mechanical transmissibility and infectivity of SnIV1 in *N. benthamiana* and in the same genotype of peppers.

In total, 25 independent studies that detected SnIV1 sequences were identified, 15 of which were gathered through the palmID search. In terms of timing, the oldest SRA dataset with SnIV1 presence was released in 2013, and is a transcriptomic study of *Medicago trunculata* root nodules from France (Roux et al. 2014), while the most recent study concerns the viromic exploration of wild *Solanum* species (Mahlanza et al. 2022) and apricots (*P. armeniaca*) (Bester and Maree 2022) in South Africa. The geographic origin of the biological samples from these studies spanned five out of the six habitable continents or 11 countries, including eight independent studies conducted in the USA. Seventeen studies involved sequencing of plant samples, including seven that involved sequencing of members of *Solanaceae* family. Of the seven studies that utilized non-plant samples, four involved the sequencing of bee species (*Apidae*). These *in silico* searches highlighted associations of SnIV1 with 12 different plant species of economic importance (*e.g.*, crop, medicinal, ornamental, or fuel feedstock), six non-crop or wild plant species, and five animal species.

### SnIV1 genomes assembled from global SRA datasets

Aside from the sequencing efforts from this study presented in Table 2, SnIV1 genomic sequences were assembled from representative SRA datasets. These are from a total of 16 studies, including 15 from palmID search hits, and one recent study identified through literature search (Table 5). In this effort, only a subset of the 80 palmID search hits were used, with the assumption that SnIV1 isolates are possibly not significantly diverse within a single study that analyzed samples collected in the same year and country, by the same research team. The datasets were from a variety of short read Illumina sequencing, with reads of between 47 and 150 nt, single or paired-end, and ranging from 3.8 to 188.3 million reads per dataset. SnIV1 genome segments were reconstructed by reference-guided assembly, with highly variable r.m.d. from of as low as 4x to as high as 77,350x. More than 96% of each genome segment was assembled, and only four out of the 63 genome segments (*i.e.,* representing 21 isolates) were not successfully reconstructed and deposited in GenBank due to partial or fragmented genome assemblies and/or of problems identified in the expected ORFs.

**TABLE 5.**
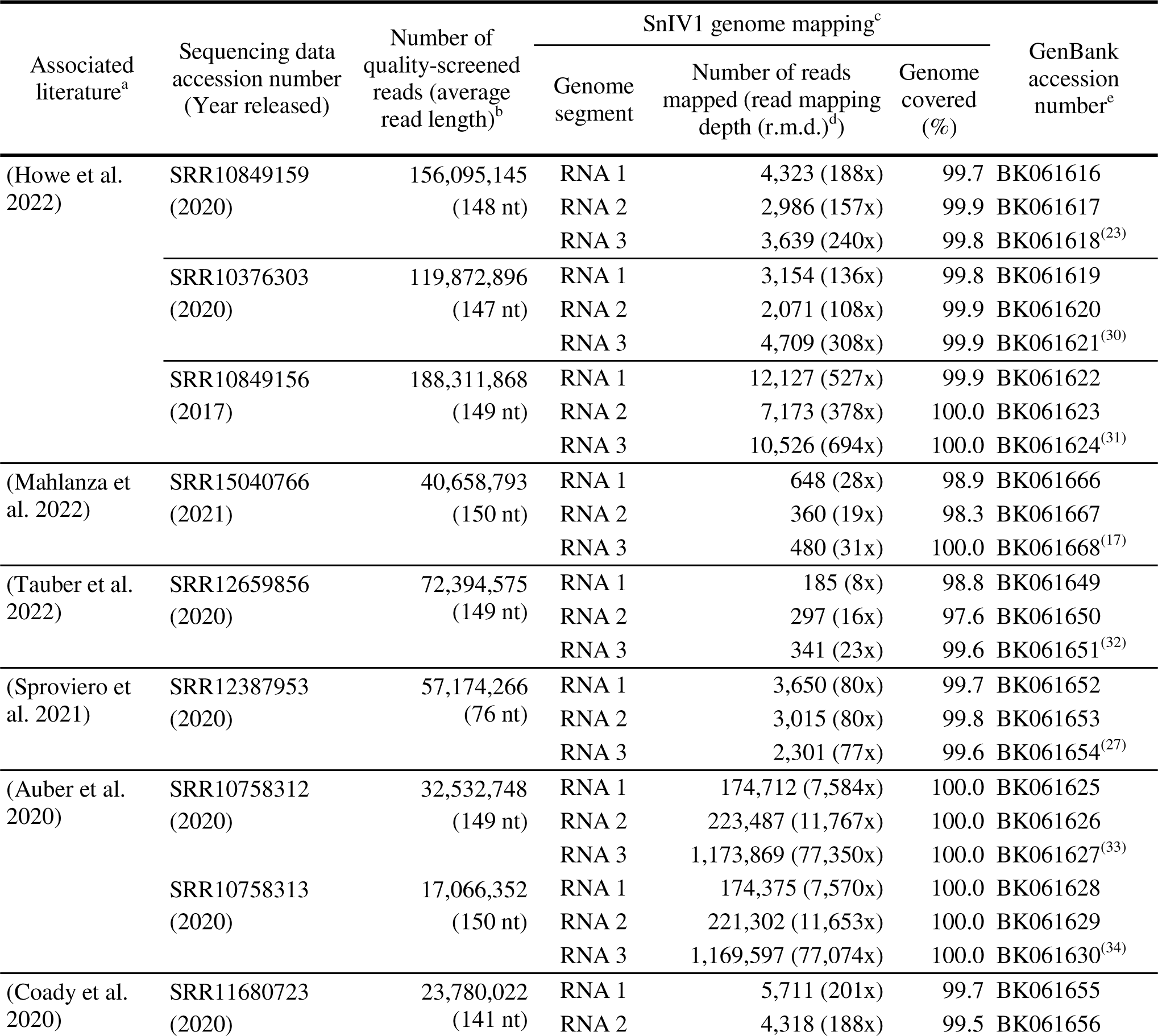
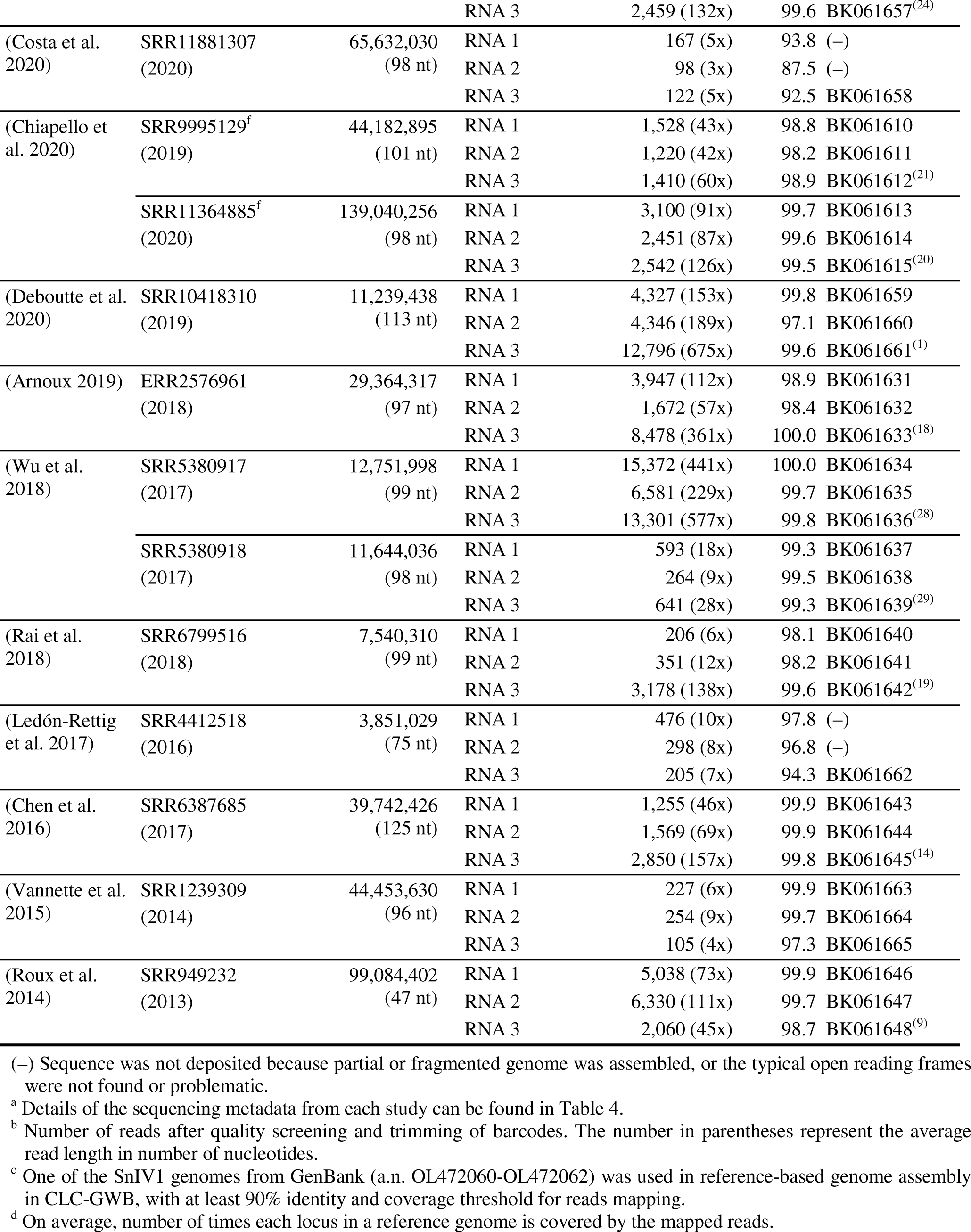
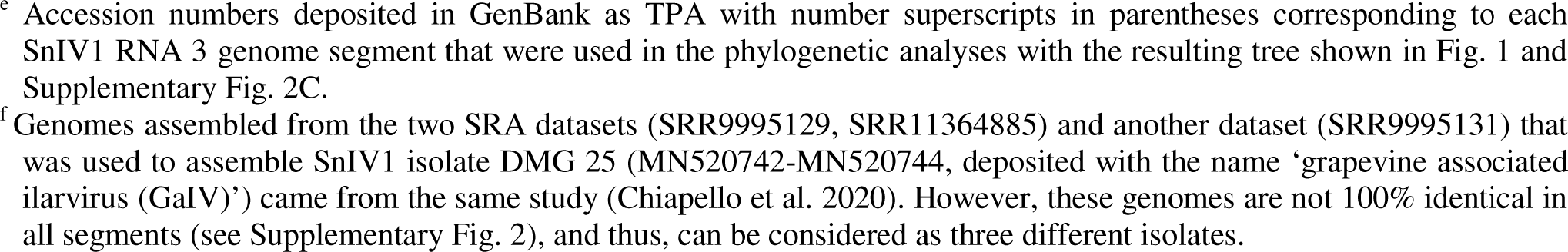
SnIV1 genome assembly from selected publicly available global sequencing data.

### Phylogenetic clustering and pairwise identity comparisons of SnIV1 global isolates

A recombination-free alignment of concatenated MP and CP ORFs (RNA 3 segment) was used to reconstruct a maximum likelihood phylogenetic tree of the global isolates of SnIV1 (Fig. 1). Four major clades or lineages were observed, including two distinct basal lineages: one comprised of four isolates from Belgium [n=2; from eggplant (a.n. OP561329) and from honeybees (a.n. BK061661)], Germany [n=1, from a *Fabaceae* weed (a.n. MN412727)], and Greece [n=1, from pepper (a.n. OP066716)], forming a monophyletic clade with 99% bootstrap support (b.s.), and the other basal lineage consisted a single isolate from Slovenia [from *Physalis* sp. (a.n. OL472062)]. Isolates from elsewhere in the world form a poorly supported clade (60% b.s.) with the Slovenian isolate as the basal lineage. The rest of the isolates formed phylogenetic clusters of mixed country or continental origin. However, there is no distinct phylogenetic clustering based on host.

**Fig. 1.**
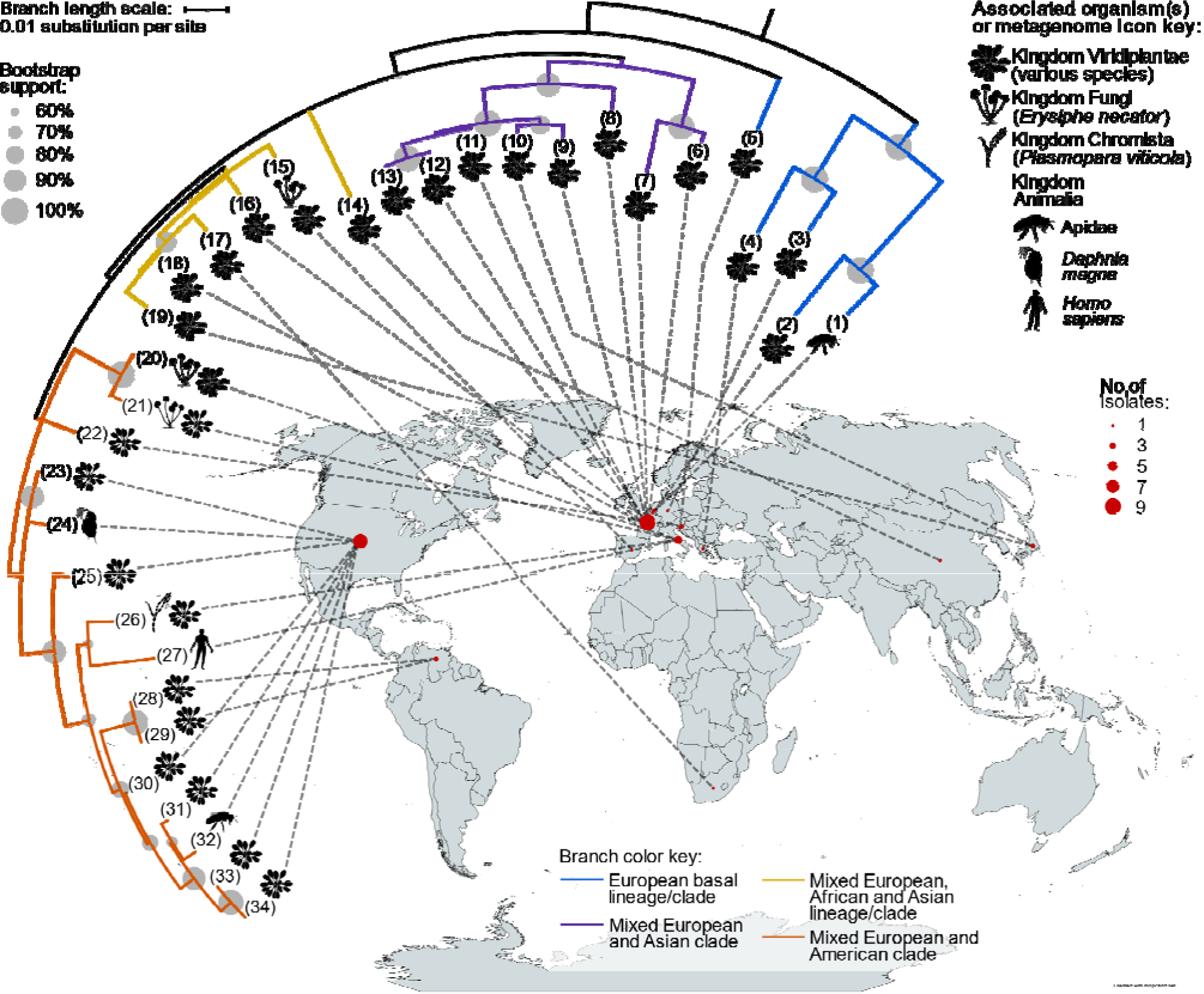
Phylogenetic clustering and host and geographical origins of 34 SnIV1 global isolates. The mid-point rooted maximum likelihood phylogenetic tree was constructed based on a multiple sequence alignment of concatenated full coding nucleotide sequences of the movement and coat proteins (RNA 3 segment). The substitution model used was Tamura 3-parameter with discrete Gamma distribution with 5 rate categories and by assuming that a certain fraction of sites is evolutionarily invariable. The tree topology shown was inferred after 1000 bootstrap replicates. Tree nodes are numbered to refer to the accession numbers of SnIV1 genome sequences indicated in Supplementary Fig. 2C, with their details in Tables 2, 4 and 5. The world map was created using MapChart (www.mapchart.net) under the CC BY-SA 4.0 license, and icons were downloaded from PhyloPic (www.phylopic.org) under the Public Domain, CC0 or CC BY-NC 3.0 licenses.

In terms of percent pairwise nt identity based on concatenated MP-CP ORFs (RNA 3), isolates of the basal European lineages are 94.5-96.6% identical to other isolates, while the rest of the isolates are 96.2-100% identical to each other (Supplementary Fig. 2). When the other genome segments are analyzed, isolates of the basal European lineage (Belgian, Greek, and German isolates only) are only 90.8-92.2% identical to the rest of the isolates using RNA 1 (methyltransferase-helicase (ORF 1a)), and 92.1-93.3% identical when using RNA 2 (RdRp and viral suppressor of RNA silencing (VSR) proteins (ORFs 2a and 2b)). Likewise, patterns of phylogenetic clustering similar to that of RNA 3 phylogenetic tree were observed in the RNA 1 and RNA 2 phylogenetic trees with the distinct basal lineage of European origin being depicted.

### Diversity of SnIV1 compared to closely related ilarviruses

Recombination-free alignments of the genome segments of SnIV1 isolates and similar alignments for isolates of its two phylogenetically closest species (TSV with wide host range and PMoV with narrow host range) were used to perform comparative diversity analyses (Fig. 2A-C). It is worthwhile to note that the number of available sequences varies among the three species. TSV has the highest number of sequences across the three genome segments, closely followed by SnIV1, while PMoV has the least number of sequences available. Inspection of π along the coding regions of each genome segment indicated that SnIV1 populations generally have lower π when compared to TSV, which has the highest overall π in any genome segment, even though a comparable number of sequences were used. This is obvious when examining genome-wide π for RNA 2. For TSV and PMoV, π values reach up to 0.27-0.37 or 27-37% probability of observing single nt differences at a single locus for any pairwise sequence comparison. However, highest π values for any segment of SnIV1 are around 0.11 only (or 11% probability of observing single nt differences). When π, as well as molecular genetic diversity (θ) and overall genetic distance were compared among the three viruses, TSV stood out with the highest values in all three segments when compared to that of both PMoV and SnIV1 (Fig. 2D).

**Fig. 2.**
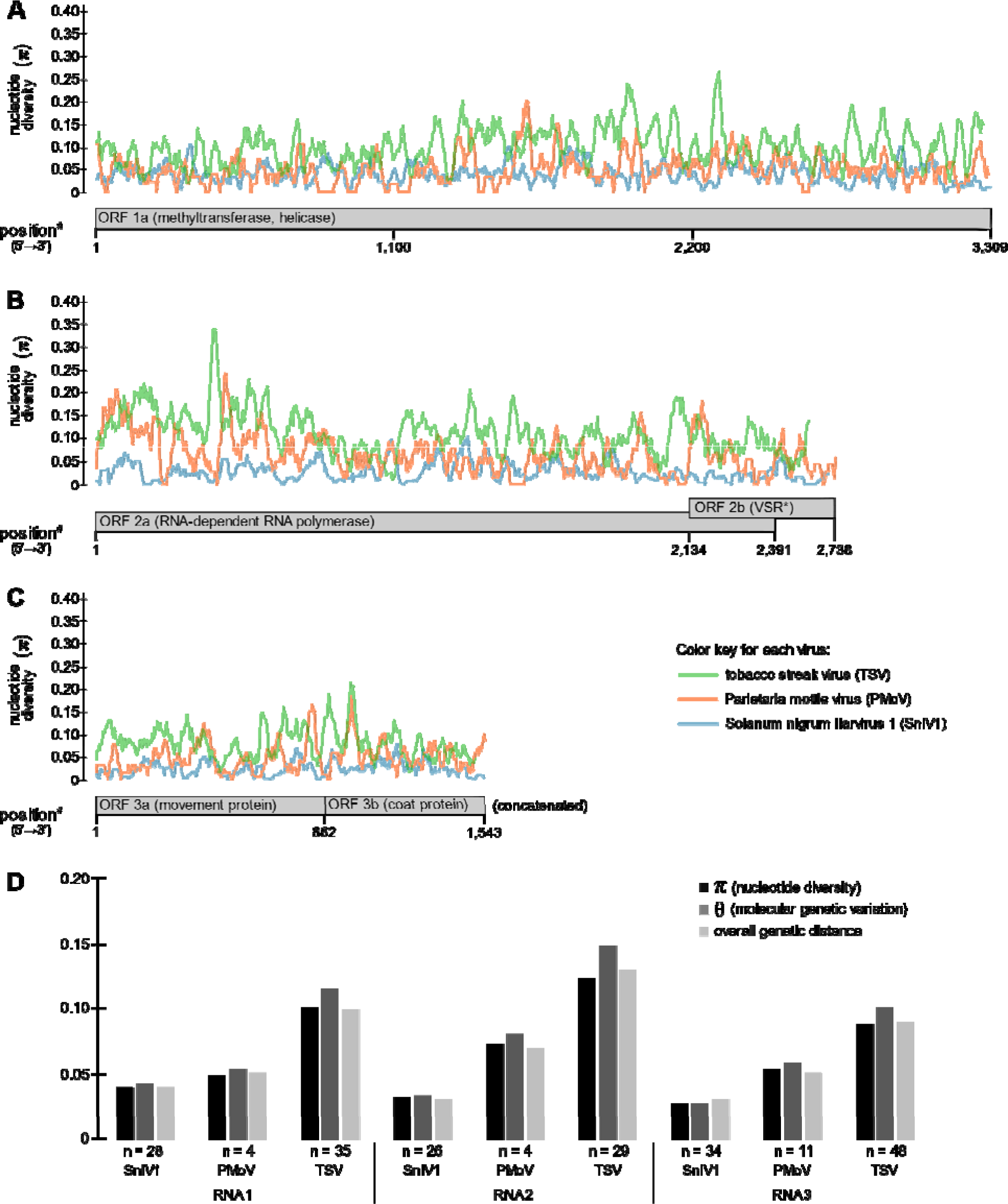
Comparative diversity analyses of global isolates of SnIV1 with those of two closely related ilarviruses, tobacco streak virus (TSV) with a known wide host range, and Parietaria mottle virus (PMoV) with a known narrow host range. **A-C**, Nucleotide diversity (π of 3) nucleotides along (A) RNA 1, (B) RNA 2, and (C) RNA calculated in window size of 30 and step 3 genome segments of TSV, PMoV, and SnIV1. **D**, measures of overall, π molecular genetic variation (θ), and overall genetic distance in all genome segments of TSV, PMoV^π^, and SnIV1. Note: ^#^position in the SnIV1 alignment. *viral suppressor of RNA silencing protein.

### Transmission and experimental host range of SnIV1

Mechanical and graft transmission assays and seed grow-out tests were performed to identify some suitable hosts and possible modes of transmission of SnIV1. Approach and chip-bud grafting experiments demonstrated that SnIV1 can be transmitted through these methods from *S. villosum* to healthy *S. nigrum* plants. Newly formed leaves of plants inoculated by both grafting methods developed symptoms such as mild vein yellowing and slight leaf crinkling (Fig. 3I,J) and tested positive for SnIV1 at 18 and 27 dpi (Table 6).

**Fig. 3.**
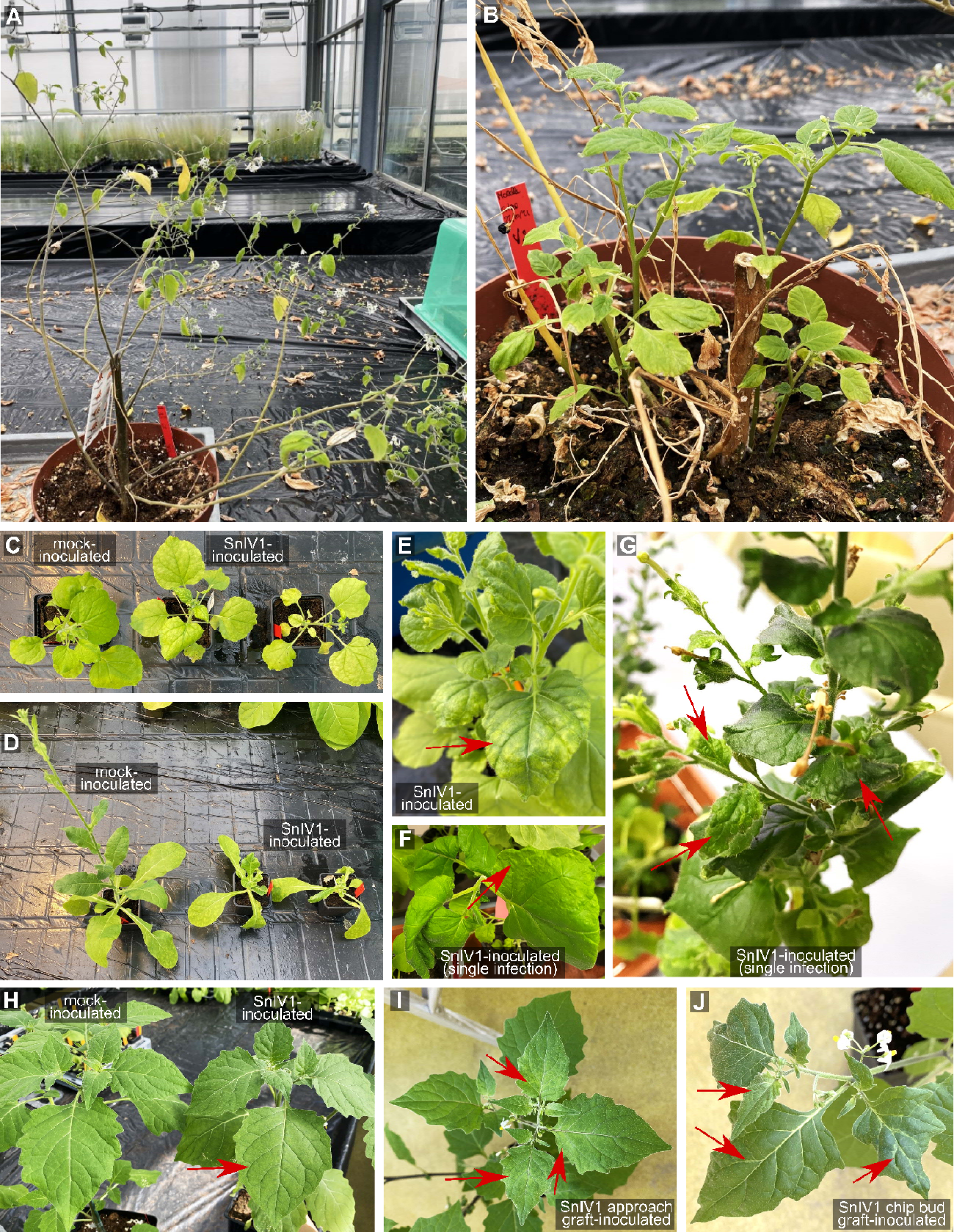
Symptoms observed in greenhouse replanted and inoculated plants. **A,B**, Two *Solanum villosum* plants collected from the field and introduced and maintained in the greenhouse. Plant shown in panel A was used as inoculum source. **C**, mock- and mechanically inoculated *Nicotiana benthamiana* plants at 28 days post-inoculation (dpi). **D**, mock- and mechanically inoculated *N. occidentalis* plants at 28 dpi. **E**, mechanically inoculated symptomatic *N. benthamiana* at 47 dpi. **F,G**, mechanically-inoculated symptomatic *N. benthamiana* at 42 dpi (F) and 105 dpi (G), that were confirmed to be singly infected b SnIV1. **H**, mock- and mechanically inoculated *Solanum nigrum* plants at 35 dpi. **I,J**, graft-inoculated *S. nigrum* plants at 35 dpi. Red arrows indicate distinct symptomatic parts of each plant shown. Inoculated plants shown in C and E were tested positive for TBRV2, but the rest of the plants shown did not undergo similar test for TBRV2.

**TABLE 6.**
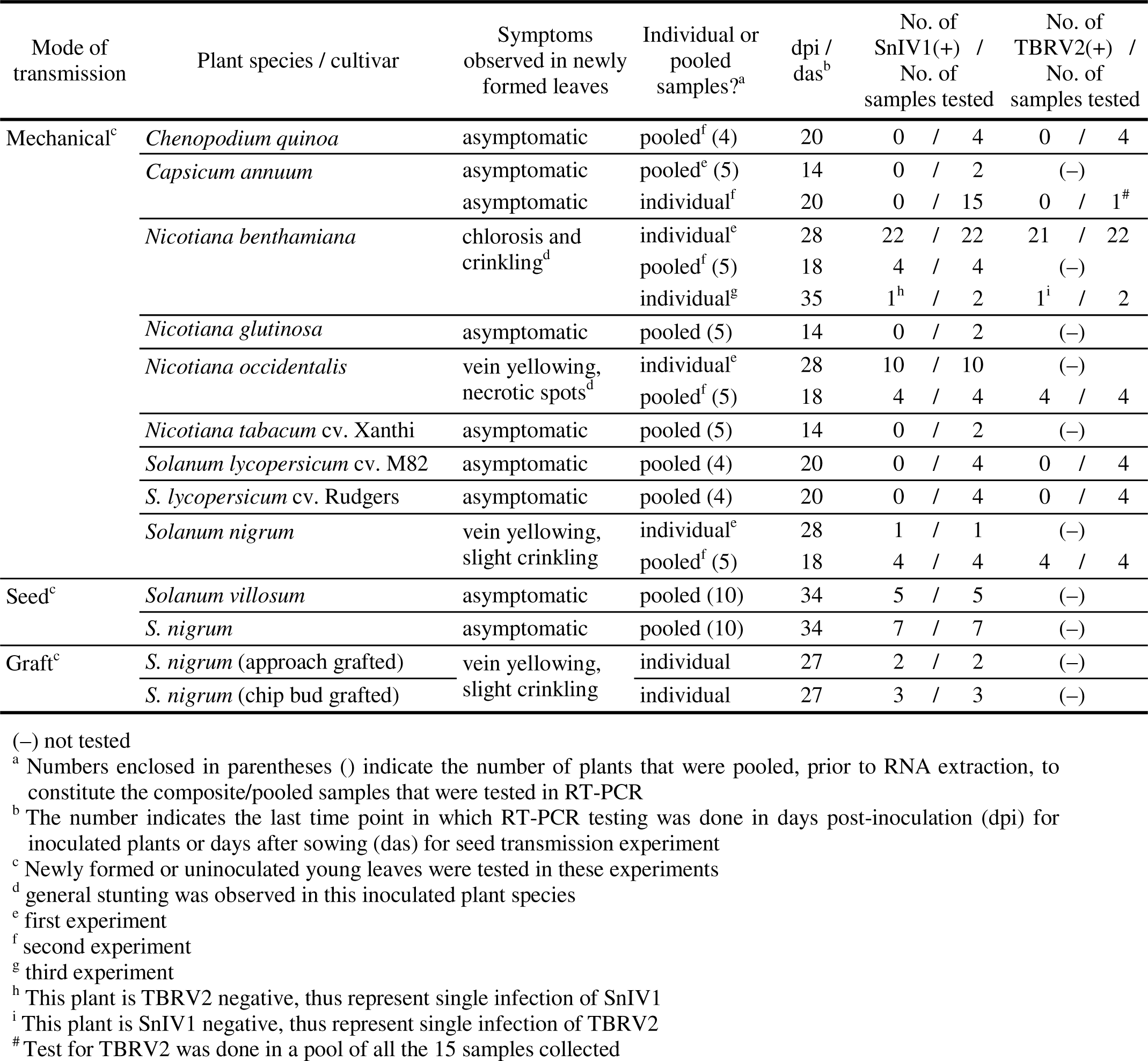
RT-PCR detection of SnIV1 and TBRV2 in plant samples from the mechanical, seed, and graft transmission experiments.

Eight plant species were mechanically inoculated, seven of which are *Solanaceae* members, including two cultivars of *S. lycopersicum.* Three solanaceous species (*N. benthamiana, N. occidentalis,* and *S. nigrum*) developed symptoms in their newly formed, uninoculated leaves and were found positive for SnIV1 when tested at 28 and 35 dpi. The rest of the mechanically inoculated plant species tested negative up until 35 dpi. The SnIV1-infected symptomatic *N. benthamiana* and *N. occidentalis* plants showed general stunting and smaller, crinkled, and chlorotic leaves compared to mock inoculated plants (Fig. 3C,D). SnIV1-infected *N. benthamiana* plants later developed more deformed leaves with interveinal yellowing (Fig. 3E). Mechanically inoculated *S. nigrum* plants showed only subtle interveinal chlorosis in their newly formed uninoculated leaves (Fig. 3H), which tested positive for SnIV1 at around 18 and 28 dpi.

SnIV1-infected *N. benthamiana, N. occidentalis,* and *S. nigrum* were later determined to be co-infected with TBRV2. Single infection of SnIV1 in a *N. benthamiana* plant was observed in a separate mechanical inoculation experiment, where the plants showed distinct bending of stems at around 28 dpi and severely crumpled leaves persisting until around 105 dpi (Fig. 3F,G). In this same experiment, a *N. benthamiana* plant singly infected with TBRV2 developed leaves with more pronounced yellowing symptoms compared to the plant singly infected with SnIV1.

Possible transmission of SnIV1 from infected seeds to newly germinated young plants was demonstrated. Seedlings of both *S. villosum* and *S. nigrum*, tested in pools of 10 leaves from different plants tested positive for SnIV1 at 34 days after sowing.

Histopathology of SnIV1-infected *Nicotiana benthamiana* and SnIV1 virion morphology.

Tissues of singly infected *N. benthamiana* collected at 49 dpi (plant shown in Fig. 3G) were examined in comparison to those of mock-inoculated plants of the same age grown under the same conditions. In healthy tissues, normal epidermal and parenchymal cells and vascular tissues were observed (Fig. 4A-C), while SnIV1-infected tissues and cells were strikingly distorted (Fig. 4D-F). While no viral particles could be observed when ultra-thin sections were examined by TEM, a high number of viral particles were readily observed on leaf dip grids with negative staining prepared using extracts from the same *N. benthamiana* leaf (Fig. 4G-I). The virions observed were spherical and measurements performed on 70 virions yielded an average diameter of 27.6 nm, with a standard deviation of 0.6 nm. Such morphological properties are consistent with those of some ilarviruses, although for some members of the genus, unstable particles or particles of slightly different sizes or with a bacilliform shape were also reported (Simkovich et al. 2021; Adams and Antoniw 2006).

**Fig. 4.**
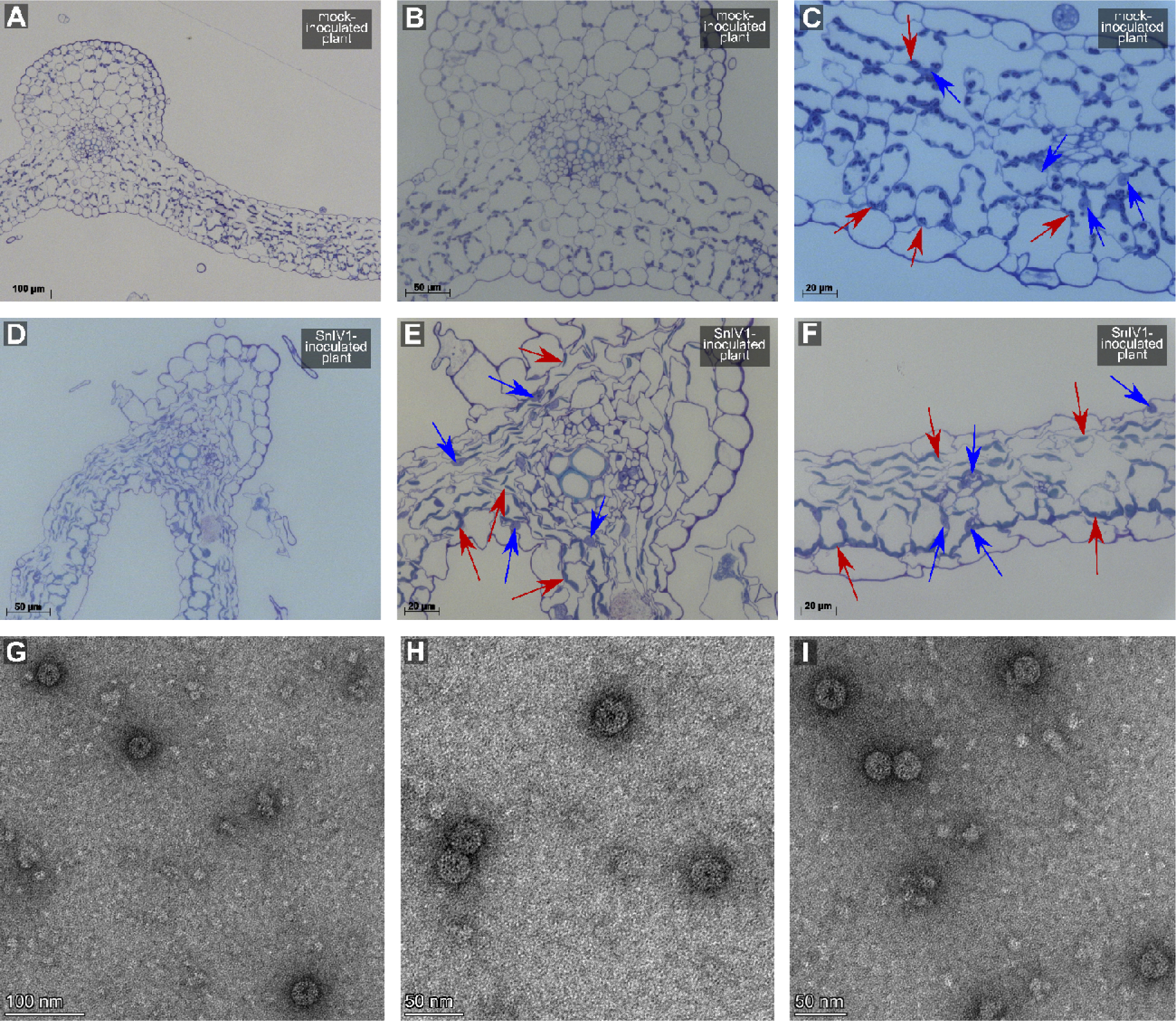
SnIV1-infected tissues and virions from mechanically inoculated *Nicotiana benthamiana* leaves. **A-C**, Thin sections of mock-inoculated *N. benthamiana* leaves for comparison that were taken at different magnifications under light microscope. **D-F**, Thin sections of SnIV1-inoculated *N. benthamiana* leaves that were taken at different magnifications under light microscope. Arrows indicate the nuclei (blu arrows) and chloroplasts (red arrows). **G-I**, Transmission electron micrograph of SnIV1 spherical virion from a crude preparation of mechanically-inoculated *N. benthami*

## DISCUSSION

### Expansion of associated hosts and geographic distribution of SnIV

HTS-based virome surveys implemented in this study uncovered association of SnIV1 with a new set of plant species from France and Belgium. Virome surveys conducted in Belgium detected SnIV1 in *S. melongena* and *S. tuberosum*, which are first detections of the virus in such species and in Belgium. This further expanded the number of *Solanaceae* species that are associated with SnIV1 that included both cultivated (*i.e.*, *S. lycopersicum, C. annuum*) or wild species (*i.e.*, *S. nigrum, S. villosum*). Transmissibility and/or infectivity of SnIV1 was only experimentally demonstrated in a limited number of solanaceous species (Orfanidou et al. 2022). Recent reports likewise associated SnIV1 with *Physalis* sp. and *S. chenopodioides* but its infectivity to them is yet to be verified (Mahlanza et al. 2022; Rivarez et al. 2022). The expanded association of solanaceous plants with SnIV1 fits very well with the results of the transmission experiments in this study, where infectivity of SnIV1 was confirmed in several solanaceous hosts.

*In silico* and literature searches implemented here likewise contributed additional information on associated hosts of SnIV1, especially those that were sampled in countries outside of Europe. Most of the plant-derived sequencing datasets with SnIV1 sequences were obtained from metagenomic or metatranscriptomic sequencing of leaf tissues of plants from different families. Interestingly, SnIV1 was detected in the metatranscriptomic sequencing of belowground parts of *Fabaceae* species (*Medicago truncatula* and *Arachis hypogaea*) (Roux et al. 2014; Chen et al. 2016) and in *Lithospermum erythrorhizon* roots (Auber et al. 2020). The HTS of *L. erythrorhizon* has the highest number of SnIV1 reads among all the datasets. In the present study, systemic infection of SnIV1 was likewise demonstrated using RT-PCR assays in different below- and above-ground parts of *S. villosum,* including roots, whole flowers, individual floral parts, and pollen. SnIV1 sequences were likewise found in a metatranscriptomic sequencing study of *Petunia* x *hybrida* (*Solanaceae*) flowers (Haselmair-Gosch et al. 2018). Collectively, such diverse host associations gave an unusual impression for a member of the *Ilarvirus* genus since its members are typically associated with a narrow range of natural hosts within one or a few families (Badillo-Vargas et al. 2016; Bratsch et al. 2019).

Moreover, SnIV1 was detected in the metatranscriptomic sequencing of insect samples from the USA and Belgium and, interestingly, in whole honeybees (*Apis mellifera*) (Deboutte et al. 2020), honeybee intestines (Tauber et al. 2022), and midgut (Vannette et al. 2015), as well as in abdomen of bumblebees (*Bombus impatiens*) (Costa et al. 2020). This information could imply the possibility of *Apidae* species harboring SnIV1 after alighting and feeding on

SnIV1-infected plants or collecting SnIV1-infected pollen from them. Such a scenario has been reported for ilarviruses associated with bee species (*Apidae*) and thrips (mostly *Thripidae*) in several studies (Bristow and Martin 1999; Sharman et al. 2015; Roberts et al. 2018; Sdoodee and Teakle 1993). Furthermore, the detection of SnIV1 sequences in bees might suggest localization of SnIV1, not only in bee integumentary parts, but also in internal parts such as the gut, however, this needs further experimental verification. The detections of SnIV1 sequences in other sequenced animal tissues such as in *Daphnia magna* and human blood are suspicious and remain to be resolved, but it is very possible that these represent contaminations that occurred at any step of the viromic experiments.

It is important to note that these findings from the viromic surveys and *in silico* search of SRA datasets do not necessarily imply that the sequenced specimens are hosts of SnlV1. Although mining of SRA datasets for viruses provided useful insights, verification of such sequence mining-derived information using another diagnostic method is difficult, thus, at least the detection of full viral genomes is recommended when analyzing metagenomic data for presence of viruses (Simmonds et al. 2017; Roux et al. 2019). Aside from true virus infection or symbiotic association with a certain organism, detection of viral sequence in an HTS dataset might result from wet lab contamination, index or barcode hopping, and cross-talk during sequencing (Lebas et al. 2022). It is likewise possible that other organisms that are the true hosts of SnIV1 could be present through an intimate (*e.g.* pathogen or symbiont) or casual association (*e.g.* as surface contaminant) with the primary sequenced sample. In such case, validation of HTS detection using a second diagnostic method and infectivity tests (if possible) are recommended (Kutnjak et al. 2021; Fox 2020).

### On the possibility of pollen as vehicle for SnIV1 spread or contamination in other plants.

Several ilarviruses and other members of family *Bromoviridae* were shown to be horizontally transmitted through pollen (Gilmer and Way 1960; George and Davidson 1963; Sdoodee and Teakle 1988; Greber et al. 1991; Mink 1993; Sdoodee and Teakle 1993; Aparicio et al. 1999; Card et al. 2007; Kawamura et al. 2014; Jaspers et al. 2015). In some cases, ilarviruses were also reported to be associated with pollen surfaces or exine (Hamilton 1977; Digiaro and Savino 1992; Fetters et al. 2022). The random directionality of transmission through pollen, whether actively or passively with the aid of arthropod vectors (Mink 1993), could result in erroneous assignation of natural host range of a virus, especially if sensitive diagnostic assays, such as HTS, are used. However, it is still an important horizontal transmission route that warrants special attention for emerging ilarviruses. For instance is PMoV, the closest phylogenetically-related virus to SnIV1, was shown to be pollen-transmitted and thus poses threat for tomato production, at least in some European countries (Aramburu et al. 2010; Aparicio et al. 2018; Parrella et al. 2020).

Based on the results of this study, it is thus hypothesized that contamination by SnIV1-infected pollen might be one of the reasons (if not the most probable) for the identification of SnIV1 in different sequencing datasets. Detection of SnIV1 sequences in 39 datasets of leaf surface or epiphyte RNA sequencing from two *Poaceae* species (*Panicum* and *Miscanthus*) (Howe et al. 2022) fits well with such a hypothesis of surface contamination by SnIV1. Thus, it is postulated that infected nearby crop or non-crop plants (most probably *Solanaceae*) could be a source of SnIV1-infected pollen that could make its way, passively or assisted by arthropod vectors, to neighboring plants. However, this hypothesis will need to be further investigated before it can be fully accepted.

Aside from plant species of the *Fabaceae*, *Rosaceae*, and *Cannabaceae*, SnIV1 was also associated with grapevines (infected with a *Erysiphe necator* and *Plasmopara viticola*) in two viromic studies (Chiapello et al. 2020, 2019), and in additional two grapevine cultivars (Ugni Blanc and Sauvignon) from France that were sequenced in this study. Yet, cuttings from two French grapevines that had positive HTS and RT-PCR detections did not shown evidence of SnlV1 presence when tested using RT-PCR several months after replanting as cuttings. This suggests either a localized or non-persistent infection in grapevine or, once again, that the first detection in field samples corresponds to surface contamination. Interestingly, another probable case of contamination is the detection of SnlV1 in *D. carota* subsp. *carota* (wild carrot). Out of 45 wild carrot populations (composite samples) (Schönegger et al., in preparation), only one showed SnlV1 reads, and this population happens to have been sampled close to the Sauvignon grapevines and SnlV1-infected *S. villosum* and *S. nigrum* plants on the INRAE Bordeaux research center.

### Patterns of variability and clustering of SnIV1 global isolates.

Relatively low diversity was observed among isolates of SnIV1, across all genome segments, when compared to closely-related ilarviruses, such as TSV for which a comparable number of sequences were available. However, this result needs to be interpreted with caution, due to the possible uneven sampling (both by number and location) of the three compared viruses. Moreover, phylogenetic analyses showed only a partial clustering of SnIV1 isolates on a geographic basis. Although there is a distinct basal clade of isolates from European countries, no distinct pattern could be observed when the time of sampling (or sequencing) or broad category of hosts (*i.e.*, plants or animals) were considered.

### Host range and other modes of transmission of SnIV1.

Two *Solanaceae* species were recently shown to be hosts of SnIV1, *C. annuum* (hybrid Arlequin F1) and *N. benthamiana* (Orfanidou et al. 2022). However, SnIV1 was not successfully mechanically transmitted to pepper plants in this study. This result might be explained by differences in inoculation procedures, inoculum source, or pepper genotypes used in the two studies. Furthermore, among eight plant species mechanically inoculated in the present study, infectivity of SnIV1 was demonstrated in three solanaceous species only: *N. occidentalis, S. nigrum,* and *N. benthamiana*. In a previous study (Ma et al. 2020), SnIV1 sequences were detected in both tomato and *S. nigrum* plants. However, in the present study, SnIV1 was successfully transmitted to *S. nigrum*, but not to the two tomatoes varieties (cv. Rudgers and M82). This suggests that association of SnIV1 with tomato in the Ma et al. (2020) study might be a result of surface contamination or, alternatively, differences in host plant or viral genotype. Nevertheless, the infectivity of SnIV1 in a genotype of pepper raises the possibility that SnIV1 could be adapting to cultivated solanaceous hosts. It is worthwhile to note that both *S. villosum* and *S. nigrum* overwinter in at least some parts of Europe and may serve as reservoir of the virus during the winter months. Overwintering of SnIV1 is likely also possible through infected seeds, since vertical transmission through seeds of these two wild species was demonstrated here.

### Summary and future perspectives.

Overall, viromic surveys, extensive sequence database exploration, and literature searches conducted in this study resulted to the expansion of knowledge on the geographic distribution, associated hosts, and diversity of SnlV1. In parallel, classical plant virology techniques facilitated the characterization of its pathobiology and possible modes of transmission. The properties of SnIV1 appear to be similar to those of other ilarviruses, but it is interestingly (and quite unexpectedly) associated much more frequently with very diverse plant samples and even with animal species mostly from *Apidae*. This raises further questions on unexplored properties of SnIV1 leading to its propensity to come up in unexpected HTS datasets. As discussed, many other ilarviruses are pollen-borne, including some infecting solanaceous species, although none have been reported in association with grapevines, despite the heavy HTS sequencing efforts performed in many laboratories around the world on this species. Although the present work facilitated progress in understanding aspects of the biology and epidemiology of SnIV1, there are still more information to be uncovered in order to fully understand this intriguing virus.

## Acknowledgments

The authors would like to thank the Sequencing Platform of Toulouse (GeT-PlaGe) for Illumina sequencing and for Marie Lefebvre for assisting in the processing of sequencing data. The authors are grateful for the assistance provided by Thierry Mauduit, Christophe Higelin and Maryam Khalili in the field, greenhouse, and sample preparation work. They are also grateful to Laure Beven for assisting in the use of dark field microscope, and Mickael Maucourt for assisting in tissue lyophilization. The authors would like to thank Varvara Maglioka and Reza M. Hajimorad for providing their SnIV1 genomic sequences ahead of release in GenBank and the publication of their papers.

## Funding information

This work mainly received funding from Horizon 2020 Marie Skłodowska-Curie Actions Innovative Training Network (MSCA-ITN) project “Innovative Network for Next Generation Training and Sequencing of Virome (INEXTVIR)” (GA 813542) under the management of the European Commission-Research Executive Agency. It was also supported by the funding from Slovenian Research Agency (ARRS) financing (P4-0165, P4-0407, J4-4553). Funding for the work in Belgium was provided by The Belgian FPS Health Food Chain Safety and Environment under Project RT18/3 SEVIPLANT. MPSR also received funding from the Balik Scientist Program (Republic Act 11035) of the Department of Science and Technology – Philippine Council for Agriculture, Aquatic, and Natural Resources Research and Development (DOST– PCAARRD), Republic of the Philippines.

## Author contributions

TC, AM, SM, MR, and DK are all involved in funding acquisition for the INEXTVIR project. TC, AM, MR, DK, and MPSR formulated and designed the study. TC supervised the study. MPSR did the majority of the experimental work, the data analyses, and wrote the first draft of the manuscript. TC contributed to sequence database mining and genome assembly. CF, AM and LSD contributed to collection and transmission experiments and RT-PCR testing of samples from France and assisted in greenhouse experiments. AP contributed in transmission experiments, nanopore sequencing and analyses, and in the transmission experiments. MTŽ did the light and transmission electron microscopy experiments. DS, SM, KDJ, and AB contributed SnIV1 sequences, and KDJ confirmed the detection of SnIV1 in Belgium by RT-PCR. All authors agreed and significantly contributed in editing the final manuscript.

## Supporting Materials and Declarations

### Supplementary files

Supplementary Figure 1. Pairwise percent identity of tomato betanucleorhabdovirus 2 (TBRV2) isolates from this study compared to known isolates from GenBank (red bold font) and other *Betanucleorhabdovirus* species.

Supplementary Figure 2. Percent pairwise identity among global SnIV1 isolates with their phylogenetic clustering, based on A, RNA 1 (1a open reading frame (ORF)), B, RNA 2 (2a and 2b overlapping ORFs), C, RNA 3 (3a and 3b concatenated ORFs).

## Data and analyses pipeline availability

Raw sequencing reads generated from this study were submitted in the NCBI Sequence Read Archive (SRA), collectively under with the BioProject accession number PRJNA874692, and in the Recherche Data Gouv platform (https://entrepot.recherche.data.gouv.fr/dataverse/inrae) with the persistent digital object identifier (doi): https://doi.org/10.57745/ZIXT4A. All SnIV1 genome sequences generated from this study were submitted to NCBI GenBank, and the accession numbers listed in Tables 2 and 6. Details of the bioinformatic pipeline used for virus genome assembly is available through this link: https://gitlab.com/ilvo/phbn-wp2-training/-/tree/master/CLC_NIB_1.

## Ethics declarations

The authors declare no competing interests.

## Notes

### Competing Interest Statement

The authors have declared no competing interest.

