## Supplementary Files for "Diversity and pathobiology of an ilarvirus unexpectedly detected in diverse host plants and in global sequencing data"

| | | | | | | | | | | |
|---|---|---|---|---|---|---|---|---|---|---|
| TBRV2 OP441766 | 100 | | | | | | | | | |
| TBRV2 OP441765 | 100 | 100 | | | | | | | | |
| TBRV2 OL472113 | **95.2** | **95.2** | 100 | | | | | | | |
| TBRV2 OL472120 | **95.2** | **95.2** | 100 | 100 | | | | | | |
| TBRV2 OL472114 | **95.2** | **95.2** | 99.7 | 99.7 | 100 | | | | | |
| TBRV2 OL472116 | **90.0** | **90.0** | 89.8 | 89.8 | 89.8 | 100 | | | | |
| TBRV1 OL472119 | 68.1 | 68.1 | 68.2 | 68.2 | 68.0 | 67.9 | 100 | | | |
| DYVV NC_028231 | 61.2 | 61.3 | 61.7 | 61.5 | 61.7 | 61.1 | 61.7 | 100 | | |
| SYVV MT185675 | 60.7 | 60.6 | 60.6 | 60.6 | 60.6 | 60.1 | 59.6 | 61.0 | 100 | |
| CdVCV1 MN273311 | 60.6 | 60.6 | 60.2 | 60.1 | 60.2 | 59.9 | 59.9 | 60.2 | 60.8 | 100 |

Heatmap scale:

- 100

- 75

< 60

**Supplementary Figure 1**. Pairwise percent identity of tomato betanucleorhabdovirus 2 (TBRV2) isolates from this study compared to known isolates from GenBank (red bold font) and other *Betanucleorhabdovirus* species. Pairwise identity values were calculated based on alignments of full length TBRV2 genomes using SDT v. 1.2. Acronyms stand for tomato betanucleorhabdovirus 1 (TBRV1), Datura yellow vein virus (DYVV), sowthistle yellow vein virus (SYVV), and Cardamom vein clearing virus 1 (CdVCV1). GenBank accession number are indicated after each virus acronym.

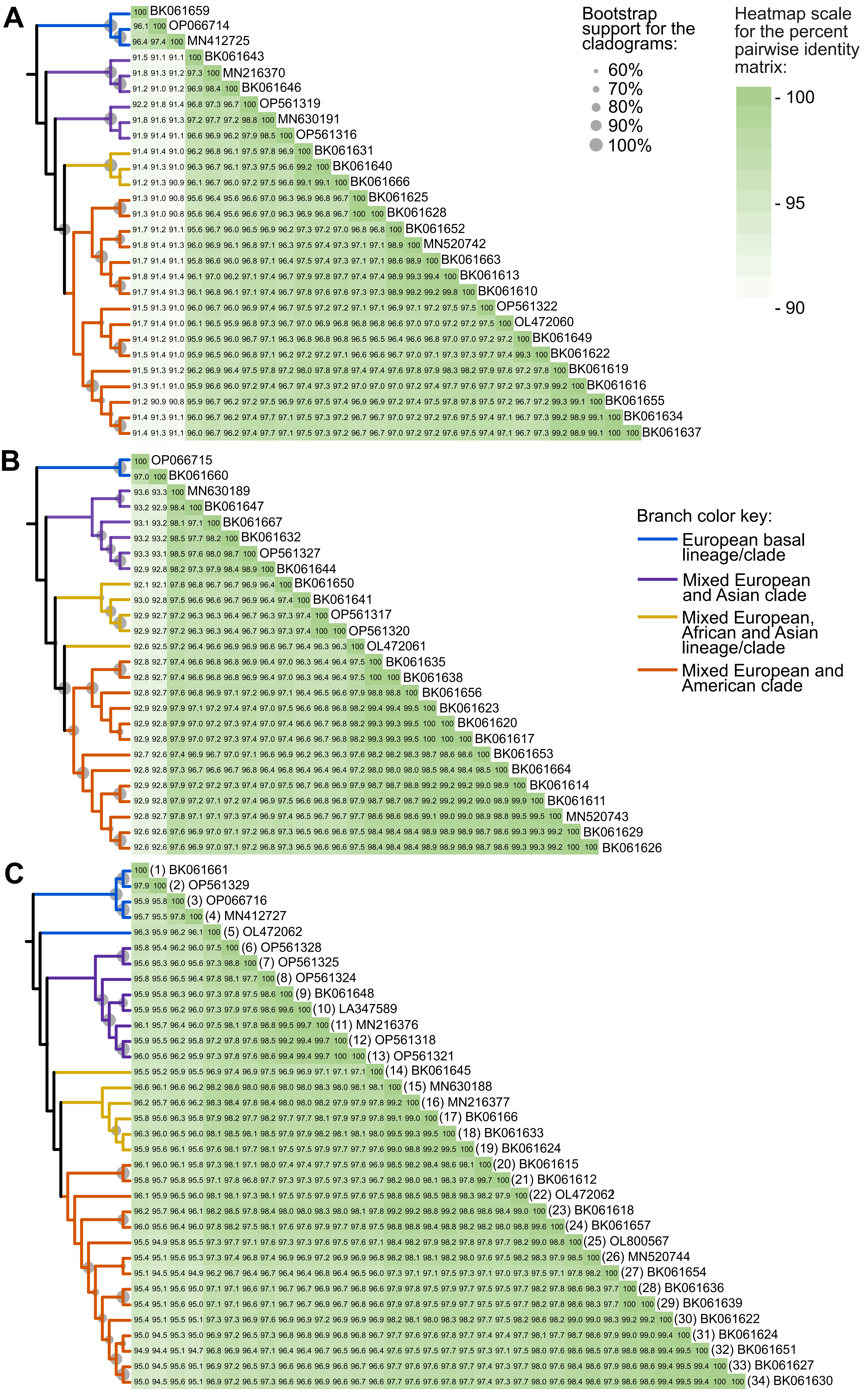


**Supplementary Figure 2.** Percent pairwise identity among global SnIV1 isolates with their phylogenetic clustering, based on **A**, RNA 1 (1a open reading frame (ORF)), **B**, RNA 2 (2a and 2b overlapping ORFs), **C**, RNA 3 (3a and 3b concatenated ORFs). The maximum likelihood phylogenetic trees were constructed based on the multiple sequence alignments of the aforementioned ORFs and were shown as consensus trees (cladograms). The substitution model used was Tamura 3-parameter with discrete Gamma distribution with 5 rate categories and by assuming that a certain fraction of sites is evolutionarily invariable. The tree topology shown as a cladogram was inferred after 1000 bootstrap replicates. The nucleotide pairwise identity was likewise calculated using the same alignments and are presented as a heatmap. Associated metadata of each sequence used in this analysis can be found in Tables 2, 4, and 5.
